# Immunoglobulin-producing AT2-like cells secrete IgA into the extracellular matrix in pulmonary fibrosis

**DOI:** 10.1101/2021.11.01.466742

**Authors:** Mengling Chen, Jing Wang, Mengqin Yuan, Min Long, Sha Wang, Wei Luo, Jie Huang, Yusi Cheng, Wei Zhang, Wei Jiang, Jie Chao

## Abstract

Pulmonary fibrosis is an interstitial lung disease that can be caused by various factors. Here, we first observed extensive IgA deposition in the extracellular matrix (ECM) of the lungs of mice with pulmonary fibrosis induced by silica inhalation. Consistent with this phenomenon, spatial transcriptomic sequencing of fresh mouse lung tissues from control mice and model mice showed that *Igha* transcripts were highly expressed in the lesion area. Single-cell RNA sequencing (scRNA-seq) and reconstruction of B cell receptor (BCR) sequences revealed a new cluster of cells with a shared BCR and high expression of genes related to immunoglobulin IgA production. Surprisingly, these clonal cells had more characteristics of AT2 (alveolar epithelial cell type 2) cells than B cells; thus, these cells were named AT2-like cells. Therefore, we propose that secretion of IgA into the ECM by AT2-like cells is an important process that occurs during lung fibrosis.

## Main

Pulmonary fibrosis is a chronic progressive disease(Noble et al., 2012). The extracellular matrix (ECM), as the framework supporting lung tissue, is mainly composed of 5 types of substances, namely, collagen, noncollagen, elastin, proteoglycan and aminoglycan(Theocharis et al., 2016). These components are affected by lung cells and can in turn act on lung cells(Mcgowan, 1992). Lung epithelial injury and apoptosis are thought to be the initiating factors of pulmonary fibrosis(Thannickal et al., 2004). To maintain the normal alveolar structure, lung epithelial progenitor cells proliferate and differentiate to replenish alveolar epithelial cells(Barkauskas et al., 2013; Liu et al., 2019). In fact, abnormal and excessive proliferation of progenitor cells fails to restore normal structure, and these cells secret various cytokines that affect fibroblasts(Katzen and Beers, 2020). In recent years, it has been reported that dysregulated repair and regeneration of epithelial cells is a driving factor of lung fibrosis(Lederer and Martinez, 2018; Parimon et al., 2020; Yao et al., 2021); however, the specific molecular mechanism by which the epithelium communicates with other cells is still unclear.

In this study, we started with lung ECM proteomics and found that IgA was the most highly deposited protein in the fibrotic lung ECM. To explore the source of IgA at the cellular and molecular levels, we applied spatial transcriptomics sequencing and single-cell sequencing to mouse lung tissue. By integrating proteomics and sequencing data, we found that AT2-like cells obviously proliferated in lungs with fibrosis induced by SiO_2_. Surprisingly, a cluster of AT2-like epithelial cells in fibrotic lungs shared the same BCR made up of *Igha*, which was exactly consistent with the immunoglobulin IgA deposited in lung ECM from fibrotic lungs. It has been reported that IgA is distributed on the surface of the mucosal membrane of the respiratory tract and promotes pulmonary fibrosis by activating fibroblasts. However, in our study, we found that a large amount of immunoglobulin IgA was deposited in fibrotic lung ECM. This is the first report in the world that epithelial cells, but not B cells, produce immunoglobulin during pulmonary fibrosis. We believe that this phenomenon is closely related to the occurrence of pulmonary fibrosis. Blocking AT2-like cells secreting IgA may be a potential way to treat pulmonary fibrosis.

## Results

### A large amount of immunoglobulin IgA is deposited in the lung ECM of mice with SiO_2_-induced fibrosis

Pneumoconiosis is a type of pulmonary fibrosis caused by inhalation of free silica. Here, we established a lung fibrosis model through intratracheal instillation of a silica suspension (50 mg/ml, 100 µl) (SiO_2_ group); the control group received normal saline (NS group). The CT and H&E staining results showed that acute inflammation was observed seven days after instillation, and collagen deposition was observed in the lungs 28 days after instillation, consistent with previous studies using this model(Liu et al., 2016; Zhou et al., 2018). At 56 days after instillation, the levels of several indicators of pulmonary function, including FEV75, IC and MMEF, were decreased (Extended Data Fig. 1a). In addition to increased lung coefficients (lung weight/mouse body weight), strip- or sheet-like high density in CT and collagen deposition were observed by Sirius red staining in the SiO_2_-56d group compared to the NS-56d group (Extended Data Fig. 1b-e). Therefore, we considered mice to have entered the fibrosis stage at 56 days after instillation of the SiO_2_ suspension.

**Fig. 1:**
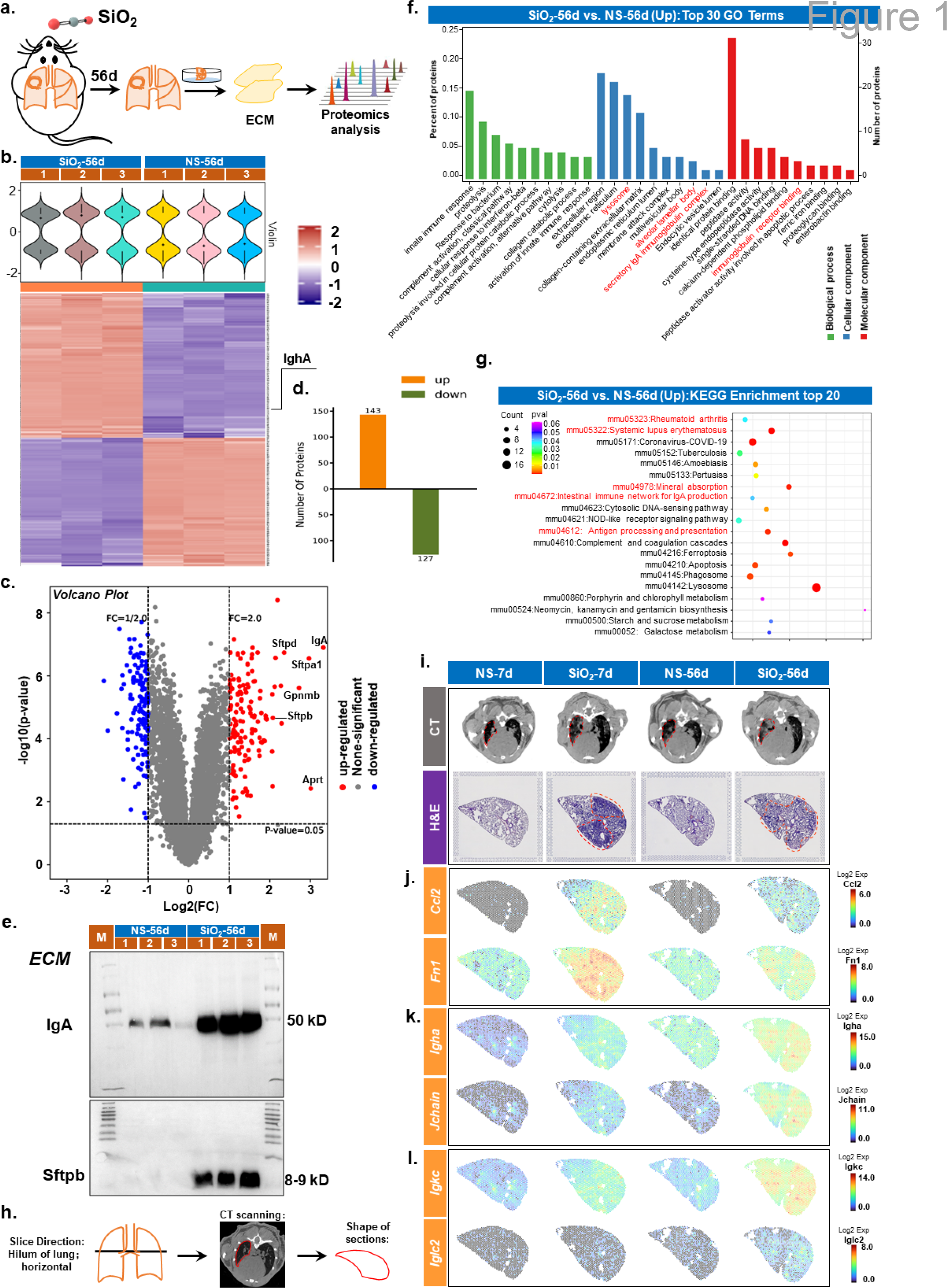
Large amount of immunoglobulin deposited in SiO_2_-56d. a, Schematic diagram of obtaining ECM. b, Heat map of differentially expressed proteins. Igha is labelled (right). c, Volcano plot of differentially expressed proteins between SiO2-56d and NS-56d. FC=2; P-value=0.05. Some markedly upregulated proteins are noted. d, Total amount of up- or downregulated proteins in SiO2-56d. d, Gene enrichment analyses of the DEGs coloured red in e. The top 30 enriched GO terms, including BP, CC and MC, are shown. Terms in red font are of particular interest. BP, biological process; CC, cellular component, MF, molecular function. f, The top 20 KEGG enrichment terms. Terms in red are of particular interest. g, Western blot of immunoglobulin IgA in ECM. h, The shape of the section derives from slicing method. Slice position: pulmonary hilum; orientation: horizontal. Because of the slicing method, it is shaped like a half moon. i, CT (top) and H&E staining (bottom) of the four samples. j-l, The corresponding gene expression in the four samples is shown. j, *Ccl2* and *Fn1* are related to inflammation; k, l, *Igha*, *Jchain*, *Igkc* and *Iglc2* are related to immunoglobulin.

To explore the ECM components of fibrotic lungs, we generated a decellularized lung matrix from NS-56d (n=3) and SiO_2_-56d (n=3) samples and analysed the protein components through mass spectrometry-based proteomics (Fig. 1a). All 6 specimens showed similar results, and the 3 NS-56d samples and the 3 SiO_2_-56d samples were most similar to each other (Extended Data Fig. 2a-b). Ultimately, we identified 143 proteins with upregulated expression and 127 proteins with downregulated expression in the SiO_2_-56d group compared to the NS-56d group (Fig. 1b-d).

**Fig. 2:**
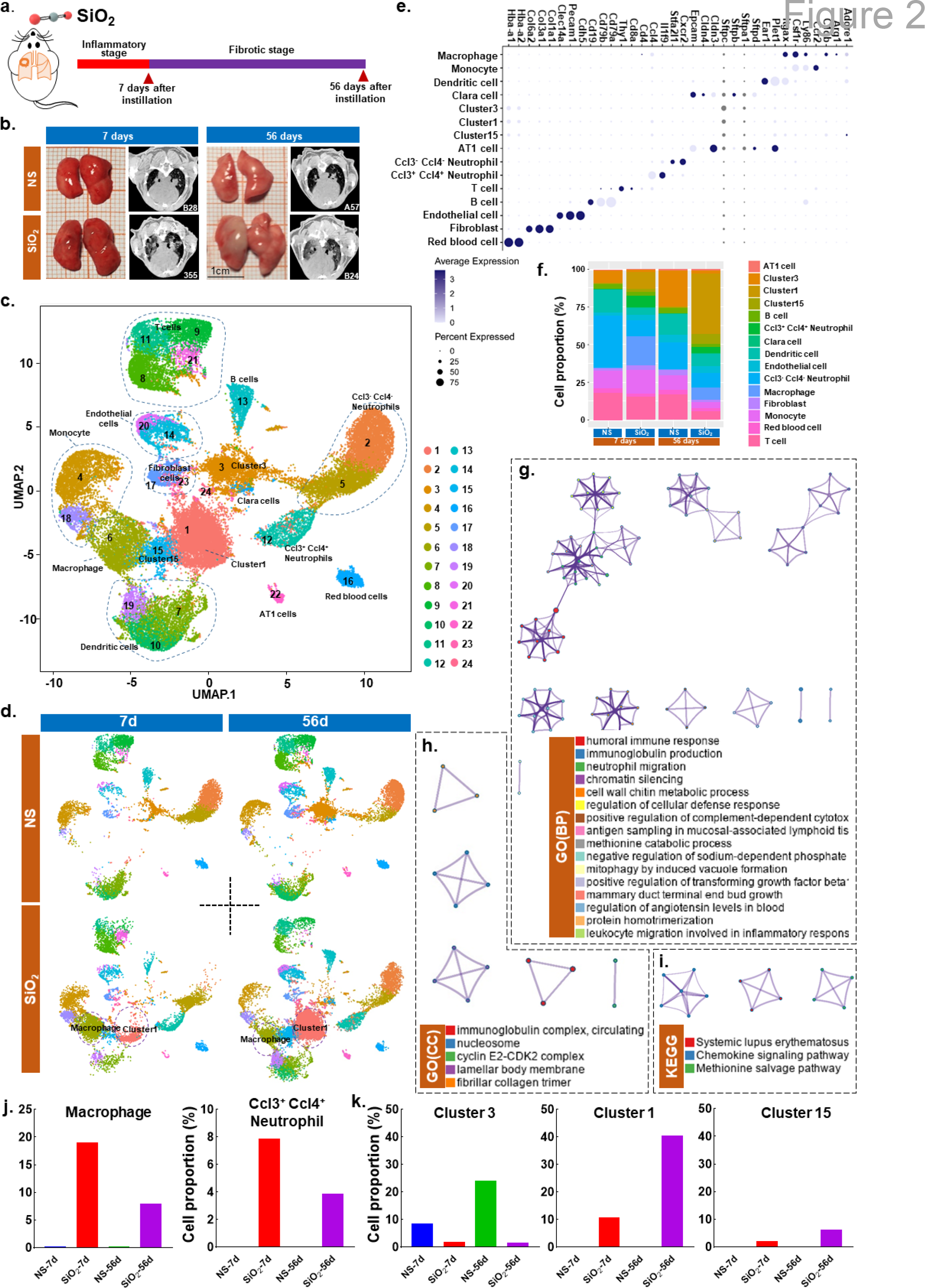
Single-cell sequencing plot of whole lung cellular transcriptional map. a, Scheme of the study design. SiO_2_-7d was used to represent the inflammation stage, and NS-7d was used as a control for SiO_2_-7d. SiO_2_-56d was used to represent the fibrosis stage, and NS-56d was used as the control for SiO_2_-56d. scRNA-seq was applied to lung cells from four samples. b, Image of fresh lung and chest CT. c, UMAP containing 43397 single cells from NS-7d, SiO_2_-7d, NS-56d, and SiO_2_-56d. A single point represents a single cell. Each of the 24 clusters is marked with a specific colour. NS-7d, instillation of normal saline for 7 days; SiO_2_-7d, instillation of silica suspension for 7 days; NS-56d, instillation of normal saline for 56 days; SiO_2_-56d, instillation of silica suspension for 56 days. d, UMAP projection of NS-7d, SiO_2_-7d, NS-56d, SiO_2_-56d samples. e, The expression of 2–3 classical cell markers of the 15 cell types assembled from the 24 clusters. f, Proportions of each cell type obtained from NS-7d, SiO_2_-7d, NS-56d, and SiO_2_-56d samples. Each cell type is labelled with a specific colour. g-i, GO and KEGG enrichment of DEGs for cluster1. Terms of interest are labelled in red. j-k, Components of related cell types at different modelling times. x axis, different samples; y axis, the proportion of this type of cells in the total number of cells from a single sample.

IgA was identified as the most highly upregulated protein (Fig. 1c). Moreover, of the top 20 proteins with upregulated expression (Fig. 1c), the surfactant proteins SFTPB, SFTPD and SFTPA are canonically thought to be secreted by mature AT2 cells(Yamamoto et al., 2017). Western blotting analysis confirmed that the IgA and SFTPB levels were substantially upregulated in the ECM of the SiO_2_-56d samples (Fig. 1e). The deposition of IgA is accompanied by the deposition of surfactant protein, suggesting that IgA may be related to AT2.

Gene Ontology (GO) analysis of upregulated proteins of SiO_2_-56d samples identified lysosome, alveolar lamellar body, secretory IgA immunoglobulin complex, and immunoglobulin receptor binding as enriched terms (Fig. 1f). The chordal graph shows that the upregulated proteins are related to extracellular regions and membrane attack complexes (Extended Data Fig. 1e). Kyoto Encyclopedia of Genes and Genomes (KEGG) analysis revealed related pathways, such as rheumatoid arthritis, systemic lupus erythematosus, and intestinal immune network for IgA production (Fig. 1g). In summary, these results indicated that lung fibrosis was closely associated with immunoglobulin production and AT2 cell activity.

To further explore the spatiotemporal characteristics of transcription in our lung fibrosis model, we performed spatial transcriptomic sequencing of four samples (NS-7d, SiO_2_-7d, NS-56d, and SiO_2_-56d). Before sequencing, all mice were scanned by computed tomography (CT) (Fig. 1i, top), and both the SiO_2_-7d and SiO_2_-56d mice exhibited hyperdense areas on their lungs, indicating that the models were successfully established. To obtain transcripts from the same structures and eliminate potential confounding by differences in anatomical position, we collected the left lung from each mouse and sliced it in the horizontal direction (Fig. 1h). Then, we observed the tissue under a microscope. If the hilum of the lung and left main bronchus were observed, the tissue section was mounted onto the spatial transcriptomics arrays. In this way, we obtained four slices (one per experimental group) with approximately the same shape and anatomical location, with the major vessels and hilum of the lung clearly visible (Fig. 1i, bottom). H&E staining showed that the alveoli of the NS-7d and NS-56d samples had a normal structure, with no infiltration of inflammatory cells (Fig. 1i, bottom). For the SiO_2_-7d sample, many inflammatory cells had accumulated, and the alveolar structures were completely destroyed (Fig. 1i). For the SiO_2_-56d sample, infiltration of inflammatory cells was still detected but was weaker than that in the SiO_2_-7d sample, and some alveoli were filled with neutrophils (Fig. 1i, bottom). This indicates that the SiO_2_-7d and SiO_2_-56d mouse models were successfully established.

After establishing the basic histology of the lung sections from each group, we performed spatial transcriptomic sequencing according to the experimental procedures recommended by 10x Genomics (details in the Methods) and obtained a total of 6778 spots (a single spot can hold cells within 60 µm in diameter). The numbers of genes and unique molecular identifiers (UMIs) detected in the tissue sections of model mice was obviously greater than that of control mice (Extended Data Fig. 3b-c). Then, we performed graph-based clustering, and we captured 11 clusters of spots (Extended Data Fig. 4a-b). The NS-7d and NS-56d samples were highly consistent in terms of gene number and clusters of spots (Extended Data Fig. 4c). The SiO_2_-7d and SiO_2_-56d samples were significantly different from their respective control samples in terms of spot clustering, and they were also significantly different from each other (Extended Data Fig. 4c). These findings indicate that mRNA expression at the same anatomical location changed substantially from the inflammatory stage (7d) to the fibrosis stage (56d).

**Fig. 3:**
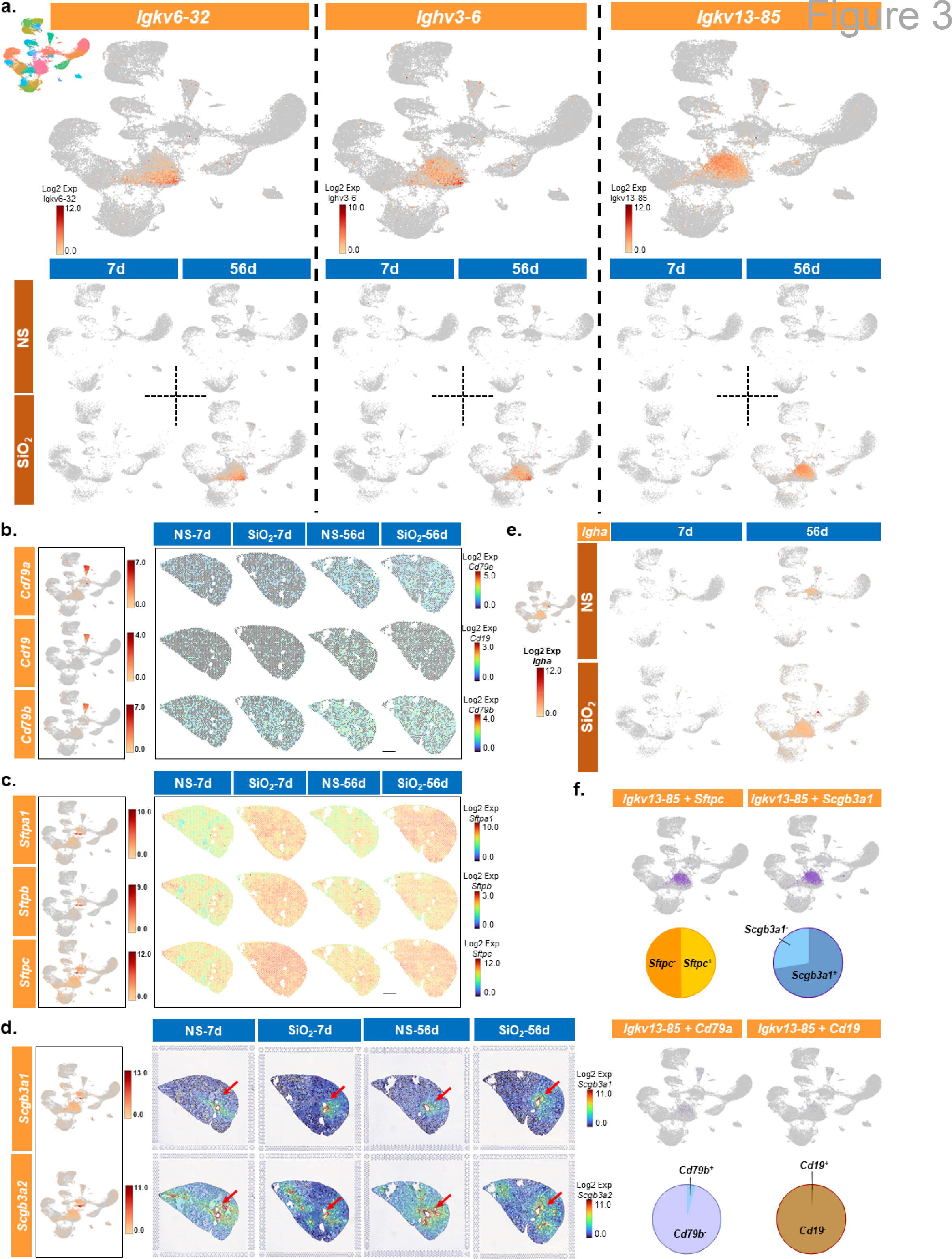
Immunoglobulin-productive cluster 1 with more characteristics of AT2 cells than B cells. a, Expression of Igkv6-32, Ighv3-6, and Igkv13-85 in the UMAP plot. The expression of each gene in four samples is displayed individually. Colour bars indicate gene expression levels. b, Expression of Cd79a, Cd19, and Cd79b in the UMAP plot and sections of four samples. c, Expression of Sftpa1, Sftpb, and Sftpc in the UMAP plot and sections of four samples. d, Expression of Scgb3a1 and Scgb3a2 on the UMAP plot and sections of four samples. Sections are merged with H&E staining image. Red arrows indicate tracheas with high expression of Scgb3a1 or Scgb3a2. e, Expression of Igha in the UMAP plot. f, UMAP plots show Igkv13-85 coexpressed with Sftpc, Scgb3a1, Cd79a, and Cd19. Fan charts show the proportions.

**Fig. 4:**
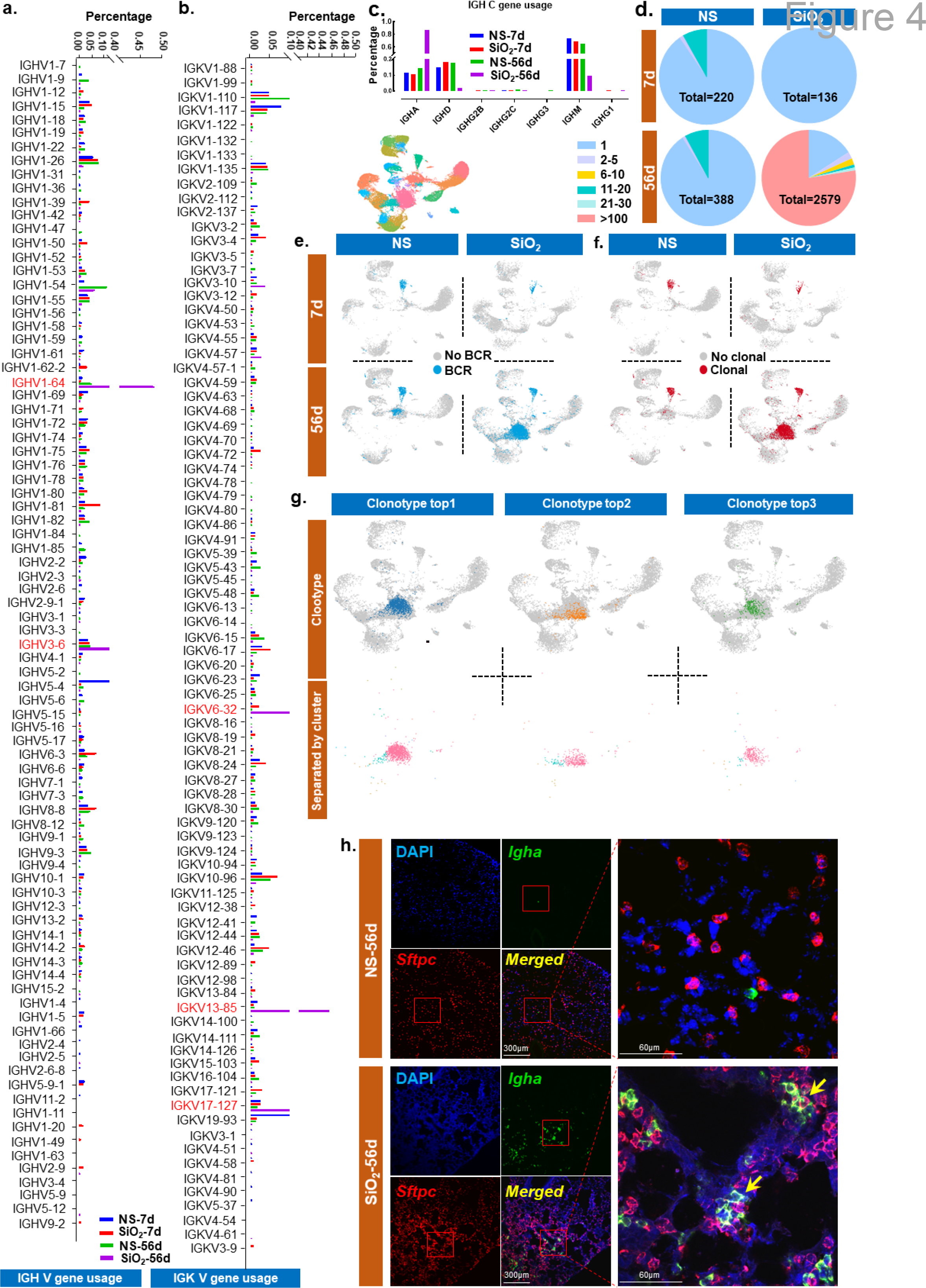
BCR clonal expansion in SiO_2_-56d. a-c, Usage of IGHV (a), IGKV (b), and IGHC (c) genes across NS-7d, SiO_2_-7d, NS-56d, and SiO_2_-56d. Different samples are shown in different colours. d, Proportion of different extents of clonotype across NS-7d, SiO2-7d, NS-56d, and SiO2-56d. e, UMAP of cells with BCR. The 24 clusters are indicated by different colours (top). f, UMAP of cells with clonotypes of BCR. g, UMAP of cells of the top 3 clonotypes (top). The cells of the top 3 clonotypes merged with the 24 clusters (bottom). Pink represents cluster 1; blue represents cluster 15. h, Igha and Sftpc detected by RNSscope. DAPI, blue; Sftpc, red; Igha, green. The regions in the red box are magnified on the right.

*Ccl2* and *Fn1*, inflammatory factors, were highly expressed in SiO_2_-7d lesions but showed decreased expression in SiO_2_-56d lesions (Fig. 1j). In terms of the expression of specific genes(Kumar et al., 2020), *Igha* and *Jchain* were highly expressed in SiO_2_-56d lesions (Fig. 1k), consistent with the observed high expression of the IgA protein (Fig. 1g). In addition, the expression of genes encoding immunoglobulin light chains, such as *Igkc* and *Iglc2*, was highly expressed in SiO_2_-56d lesions (Fig. 1l). Combined with proteomic findings for the lung ECM, these results indicated a strong relationship between immunoglobulin expression IgA and lung fibrosis.

### Single-cell sequencing detected highly amplified AT2-like cells in the SiO_2_-56d sample

To identify the cellular origin of IgA, we obtained whole lungs from four mice (NS-7d, SiO_2_-7d, NS-56d, and SiO_2_-56d, model time and groups were consistent with those used for spatial transcriptomic sequencing) and performed scRNA-seq. All mice were scanned by CT prior to tissue collection to confirm effective modelling (Fig. 2b). After scRNA-seq, the numbers of UMIs and genes were evaluated, and a total of 43,397 cells were obtained for data analysis (Extended Data Fig. 5a-c). We normalized the transcriptomic data from the four groups and then applied graph-based clustering to the combined cells from all groups to analyse and compare the samples, which yielded 24 cell clusters (Fig. 2c). A heat map of the top 20 genes expressed in each cluster is shown in Extended Data Fig. 5d. We then annotated the cell type for each cluster according to canonical cell markers from Cell Marker(Zhang et al., 2019) and assigned these 24 clusters to 15 types of cells (Fig. 2e), including endothelial cells, monocytes, macrophages, dendritic cells, B cells, T cells, Ccl3^-^ Ccl4^-^ neutrophils, Ccl3^+^ Ccl4^+^ neutrophils, fibroblasts, AT2-like cells (cluster 1, cluster 3, cluster 15), red blood cells, and AT1 cells. The marker genes of the different cell types are shown in the uniform manifold approximation and projection (UMAP) diagram (Extended Data Fig. 5e).

**Fig. 5:**
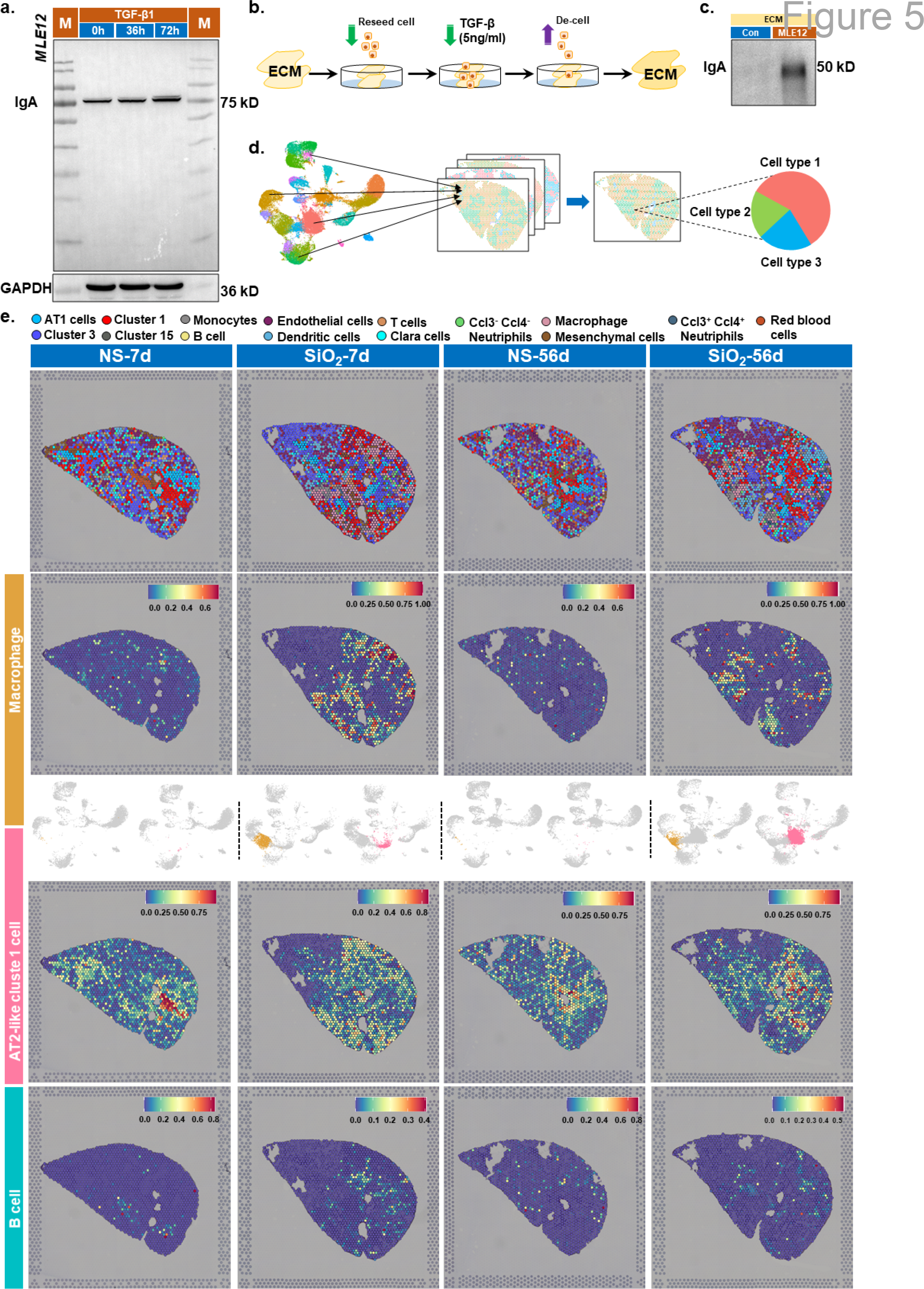
Verification through in vitro experiment and data conjoint analysis. a, Western blot of immunoglobulin IgA in mouse MLE-12 cells. b, Schematic diagram for reseeding cells. c, Western blot of immunoglobulin IgA in the ECM after reseeding cells. d, Diagram for the conjoint analysis of scRNA-seq and spatial RNA-seq. e, Predicted distribution of 15 types of cells in the four samples. Different colours represent corresponding cell types. Heat maps show the scores of macrophages, AT2-like cluster 1 and B cells on each spot. Macrophages and AT2-like cluster 1 cells across the four samples is shown in UMAP.

To explore the cell compositions of the different samples, we calculated the relative percentages of the main 15 cell types in the four samples (Fig. 2f, Extended Data Fig. 5f). The relative percentage of macrophages in the SiO_2_-7d sample was obviously increased compared to NS-7d, representing up to 18% of the total number of cells, and the percentage remained higher than that in the NS-56d sample, even in the SiO_2_-56 sample (Fig. 2j), indicating that macrophages were major participants in the inflammatory stage and were still present in the fibrotic stage. GO and KEGG analyses of differentially expressed genes (DEGs) of macrophages identified genes classically involved in the development of silicosis (Tan and Chen, 2021) (Extended Data Fig. 6). Ccl3^+^ Ccl4^+^ Neutrophils, which are often involved in various inflammatory reactions induced by bacteria, viruses, and foreign objects(Liew and Kubes, 2019), were activated upon stimulation with silica after 7 days and were detected until day 56 (Fig. 2j).

Because cells of clusters 1, 3, and 15 expressed *Sftpc*, *Sftpb* and *Sftpa1*, we identified the three clusters as AT2-like cells. The proportion of AT2-like clusters 1 and 15 both showed an increase, but cluster 15 had far fewer cells than cluster 1 (Fig. 2k). Interestingly, the proportion of AT2-like cluster 1 cells in the SiO_2_-56d sample was notably increased, reaching 40% (Fig. 2k). GO and KEGG analyses of up-regulated genes (Fc>2, p<0.05) of AT2-like cluster 1 cells indicated that they were enriched in immune-related terms, such as systemic lupus erythematosus; immunoglobulin complex, circulating; humoural immune response; and immunoglobulin production (Fig. 2g-i). Thus, we hypothesized that AT2-like cluster 1 plays an important role during the fibrosis stage.

### Immunoglobulin-producing AT2-like cluster 1 cells have more characteristics of AT2 cells than B cells

To gain further insight into the function of cluster 1, we focused on the DEGs in this cluster compared with all other cell clusters. We found that *Igkv13-85*, *Ighv3-6* and *Igkv6-32* were the top 3 DEGs in AT2-like cell cluster 1 (Fig. 3a). All three belong to the V gene group, with *Igkv13-85* and *Igkv6-32* encoding the mouse immunoglobulin light chain and *Ighv3-6* encoding the heavy chain(Babbage et al., 2006). Furthermore, all three genes were expressed at high levels in the SiO_2_-56d sample and were mainly expressed in cluster 1 cells (Fig. 3a). Moreover, *Igha* was verified to be highly expressed in cluster 1 in this sample (Fig. 3e), consistent with the observed high levels of IgA deposition (Fig. 1g).

These genes code for immunoglobulin and are thought to be expressed by B cells. To ensure that cluster 1 was not B cells, we analysed the expression of several B cell marker gene distributions in all the clusters by Loupe Brower 4.0. B cell marker genes such as *Cd79a*, *Cd79b* and *Cd19* were relatively highly expressed in the noted B cells in Fig. 2c (Fig. 3b left). However, in the sections, no visible differences in the expression of these genes were observed between the SiO_2_-7d and NS-56d or SiO_2_-56d and NS-56d samples (Fig. 3b right). This phenomenon indicated that B cells did not play a major role in the disease.

In contrast, genes encoding AT2 markers such as *Sftpc, S*ftpa1 and *Sftpb*(Han et al., 2018; Jacob et al., 2017) were relatively highly expressed in cluster 1 (Fig. 3c left) and showed significantly upregulated expression in the SiO_2_-7d and SiO_2_-56d sections (Fig. 3c right). *Scgb3a1 and Scgb3a2*, markers of secretory epithelial or Clara cells(Naizhen et al., 2019; Reynolds et al., 2002) (the progenitor cell of AT2), were both expressed in AT2-like cluster 1 cells (Fig. 3 d left). Spatially, they were expressed in the trachea (Fig. 3 d right). As these five genes (*S*ftpa1, *Sftpb*, *Sftpc*, *Scgb3a1* and *Scgb3a2*) showed relatively higher expression in cluster 1, our findings suggested that cluster 1 cells originated from bronchial epithelial cells.

Furthermore, of the five genes, *S*ftpa1, *Sftpb* and *Sftpc* were more likely to be activated in the SiO_2_-56d sample than in the NS-56d sample (Fig. 3c).

Simultaneous examination of the expression of *Igkv13-85* and cell markers of AT2 or Clara cells revealed that more than half of the *Igkv13-85^+^* cells coexpressed *Sftpc* (AT2 cell marker) or *Scgb3a1* (Clara cell marker), but few coexpressed *Cd79a* or *Cd19* (Fig. 3f). This result strongly suggested that these *Igkv13-85^+^* cells were likely not derived from B cells. Based on the analysis results, we speculated that cluster 1 represents a specialized type of AT2 cell (alveolar epithelial cell type 2). In other words, silica inhalation could trigger the transformation of normal AT2 cells into *Igkv13-85^+^* AT2-like cells, which can transcribe BCR genes.

### BCR clonal expansion in the SiO_2_-56d sample

Because we observed the differential expression of the gene encoding the immunoglobulin light chain in the SiO_2_-56d sample, we wanted to further explore the usage of the V(D)J and C genes of BCR in the four samples. We reconstructed BCR sequences from scRNA sequencing. Across the four groups, several V genes, including *Ighv1-64*, *Igkv13-85*, *Ighv3-6*, *Igkv6-32* and *Igkv17-127*, were found at notably higher frequencies in the SiO_2_-56d sample (Fig. 4a-b). Among these five genes, *Igkv13-85*, *Ighv3-6*, and *Igkv6-32* were categorized as highly expressed DEGs in AT2-like cell cluster 1 (Fig. 3a). *Ighv1-64* and *Igkv17-127* were also mainly expressed in AT2-like cell cluster 1(Extended Data Fig. 7a-b). Among IGH C genes, the expression of *Igha* was substantially increased in the SiO_2_-56d sample compared with the other three samples (Fig. 4c), which also indirectly led to a decline in the proportion of *Ighm* and *Ighd* usage. There was no obvious difference in the usage of other BCR genes across the four samples (Extended Data Fig. 7c-f).

On the basis of traditional immunology theory, we believe that clonal expansion is closely related to disease progression. According to an analysis of the top 30 clonotypes from each sample, we found no identical clonal expansions among the four samples (Extended Data Fig. 8a-d). A heat map composed of different V-J gene pairing frequencies showed 9 kinds of V-J gene combinations in the SiO_2_-56d sample in clonal populations that had expanded to over 100 cells (Extended Data Fig. 9a-d). In the SiO_2_-56d sample, clonotypes comprising over 100 cells constituted more than 75% of the total clonal cell population (Fig. 4d), and approximately 75% of the total clonal cells came from AT2-like cluster 1 (Extended Data Fig. 7h). Combined with the scRNA sequencing information, these results showed that BCR genes were expressed not only in B cells but also in AT2-like cluster 1 cells. In fact, more BCR^+^ cells were identified as AT2-like cluster 1 cells. (Fig. 4e). As with BCR, at least half of the clonal cells were identified as AT2-like cluster 1 cells through scRNA barcodes (Fig. 4f). These clones produce IgA with the progression of pulmonary fibrosis. To determine which cluster of cells the clones came from, we merged the barcodes of the top 3 clonal cell populations with barcodes from single-cell sequencing. Most of these clonal cells were identified as AT2-like cluster 1 cells (Fig. 4g). Compared with B cells, top 1 clonal cells highly expressed *Sftpc* and *Sbgb3a1* (Extended Data Fig. 7i).

The use of the V gene varies among different individuals. As expected, *Ighv1-64*, *Igkv13-85* and *Igkv6-32* were barely expressed in the sections used for spatial transcriptomic sequencing (Extended Data Fig. 10a). We next focused on *Igha*, a gene encoding the constant region of the immunoglobulin heavy chain that is crucial for determining the immunoglobulin isotype(Kumar *et al*., 2020). As shown, this gene was coexpressed with *Sftpc* (Extended Data Fig. 10b). Moreover, the expression of *Igha* in the SiO_2_-56d sample was much higher than that in the SiO_2_-7d sample, consistent with the expansion of IgA-related clonotypes in the SiO_2_-56d sample (Extended Data Fig. 7g). Therefore, the increased frequency of *Igha* use is common in pathological conditions. The IgA-expressing AT2-like cell population might be important in pulmonary fibrosis.

To verify the presence of IgA-expressing AT2-like cells in fibrotic lung tissue, we used RNAScope, a new technology that can simultaneously detect multiple target RNAs in situ. Probes for detecting *Igha* and *Sftpc* transcripts were applied. Cells containing *Sftpc* transcripts were scattered throughout the NS-56d lung sections, and these cells were normally located in the alveolar region (Fig. 4h, Extended Data Fig. 10 c). In the SiO_2_-56d sections, *Sftpc^+^* cells tended to be crowded together. In the NS-56d sections, we found only a few cells containing *Igha* transcripts. Additionally, *Igha* positivity was substantially increased in the SiO_2_-56d sections, and the *Igha* transcripts were colocalized with the *Sftpc* transcripts within the same cells, that is, AT2-like cluster 1 cells.

Based on the phenomena described above, we concluded that the increased *Igha* transcripts in the SiO_2_-56d group were derived from AT2-like cluster 1 cells.

### TCR is not amplified in the SiO_2_-7d or SiO_2_-56d sample compared with the NS-7d or NS-56d sample, respectively

T cells are primarily responsible for cellular immunity, as opposed to humoral immunity. To evaluate the role of T cell immunity in lung fibrosis, we rebuilt TCR sequences for the NS-7d, SiO_2_-7d, NS-56d, and SiO_2_-56d samples from scRNA-seq. There were no notable differences in V(D)J and C gene usage among the four samples (Extended Data Fig. 11a-f). The amino acid sequences of the top 10 complementarity determining regions (CDR3) in the four samples were different, without a specific pattern or shared CDR sequences (Extended Data Fig. 11g). The extent of clonal expansion in the four samples showed that more than 80% of the clonotypes of each sample were represented by only one cell (Extended Data Fig. 11h), and the clonotype with the greatest expansion represented no more than 3% of all cells. Then, to explore the similarities and differences in clonotypes among the four samples, we analysed the top 30 clonotypes of each sample. Each sample had a unique preferred clonotype, and no clonotype was shared among the four samples (Extended Data Fig. 12a).

Then, we used Loupe (version 4.1.0, 10x Genomics) to integrate barcodes of scRNA sequences and TCRs. Briefly, cells harbouring TCRs or paired clonotypes were recognized as T cells in each sample, and there were no notable differences across the four samples (Extended Data Fig. 12b-c). Together, these findings indicate that there were no notable differences among the samples, suggesting that T cells were not engaged in lung fibrosis.

### Immunoglobulin IgA was increased in mouse lung epithelium under TGF-β treatment

TGF-β is a known strong fibrotic factor that is commonly used to stimulate cells to establish in vitro fibrosis models. Here, we cultured MLE-12, a mouse lung epithelial cell line with the characteristics of AT2, and treated it with TGF-β for different times. Then, we evaluated IgA by Western blotting. Using an anti-IgA antibody, we detected IgA, a band at 75 kDa, exactly equal to the total molecular weight of a light chain and a heavy chain of IgA(Snoeck et al., 2006). The band became darker after the cells were treated with TGF-β for 72 hours. (Fig. 5a). To verify that AT2 cells can secrete IgA into the ECM, we reseeded MLE-12 cells on lung ECM obtained from healthy mouse lungs, followed by treatment with TGF-β (Fig. 5b). Western blotting analysis showed that more IgA was detected in the ECM cultured with MLE12 cells and TGF-β than in pure lung ECM (Fig. 5c). This phenomenon indicates that the lung epithelium secretes IgA to the lung ECM.

### Combined analysis of scRNA-seq and spatial transcriptomics showed that more AT2-like cluster 1 cells were accumulated in the SiO_2_-56d sample

To further investigate the location of cluster 1 cells in the lungs in relation to fibrotic lesions, we used Seurat V3.2 to integrate our datasets based on anchor features and predicted the main cell types included in the spatial transcriptional spots in the four sections (NS-7d, SiO_2_-7d, NS-56d, SiO_2_-56d) (Fig. 5d). All 15 cell types (Fig. 1e) identified by single-cell sequencing were predicted in the four sections. All types of cells were scattered across the four groups of samples without any obvious pattern (Fig. 5e), unlike the case in the brain and kidney, in which specific cell clusters were associated with particular anatomical locations(Asp et al., 2019; Ferreira et al., 2021). As expected, macrophages were more densely clustered in the SiO_2_-7d sample than in the NS-7d sample (Fig. 5e), consistent with their increased numbers observed by scRNA-seq (Fig. 2k). Similarly, AT2-like cluster 1 cells were found at a greater density in lesions of the SiO_2_-56d sample than in those of the NS-56d sample (Fig. 5e). Notably, there was no special pattern of distribution for B cells in the SiO_2_-56d sample (Extended Data Fig. 13a). From a spatial perspective, this result verified that macrophages participated in silicosis, especially in the early stage and that BCR^+^ AT2-like cells tended to play an important role in the fibrosis stage.

## Discussion

Fibroblasts are generally believed to play an important role in pulmonary fibrosis(Thannickal *et al*., 2004). Although drugs that block fibroblasts can slow the decline in lung function in patients with pulmonary fibrosis, they cannot reduce the mortality of patients. Therefore, there is still much work to do to identify the important events and molecular mechanisms in pulmonary fibrosis to provide targets for pulmonary fibrosis treatments. Our study found that AT2-like cells proliferated and secreted IgA into the lung ECM. These cells account for much more than other lung cells in the total number of cells, and the activation of these cells greatly changes the composition of lung ECM.

IgA is usually distributed on the mucosal surface of the digestive tract and respiratory tract and is secreted by B cells. In our pulmonary fibrosis model, IgA is deposited in large amounts in the lung ECM, which is the first report in the world thus far. In fact, pulmonary fibrosis disease is clinically associated with autoimmune diseases such sarcoidosis and systemic sclerosis(Turnerstokes et al., 1990). Studies have shown that IgA in fibrotic lungs can activate fibroblasts, promote their transdifferentiation into myofibroblasts and secrete collagen. Our findings provide a basis for fine contact between IgA and fibroblasts.

Lung alveolar progenitor cells can self-renew, proliferate and replenish damaged alveolar epithelium(Juul et al., 2020). When they proliferate abnormally, they cause alveolar structural disorders(Parimon *et al*., 2020; Yao *et al*., 2021). Three kinds of lung progenitor cells have been reported. In our study, AT2-like cell cluster 1 was more similar to Sftpc^+^ Scgb3a1^+^ lung progenitor cells, and they secreted IgA to the ECM. Although immunoglobulins are thought to be derived from B cells, studies have shown that in lung cancer, epithelial cells can also secrete immunoglobulin IgG(Qiu et al., 2003). This is the first discovery that epithelial cells produce immunoglobulins in noncancer diseases.

It has been suggested that idiopathic pulmonary fibrosis will be reclassified in the next 10 years(Wells et al., 2018). With this in mind, the findings in this study inspire us to investigate other types of pulmonary fibrosis and classify them according to the presence or absence of IgA deposition. Targeting the ablation of overproliferating AT2-like cells or blocking the production of IgA may become a potential target for the treatment of pulmonary fibrosis.

## Extended data

### Ethics

All animal experiments were approved by the Laboratory Animal Care and Use Committee of Southeast University. All procedures were conducted in accordance with the Declaration of Helsinki.

### Mice

Male C57BL/6 mice were purchased from Hangzhou Ziyuan Experimental Animal Co., Ltd. Three to five mice were raised in a cage at a stable temperature (22 °C ± 2 °C) and humidity (40% ± 10%) with a 12-h light–dark cycle. Mice were allowed free access to food and water at specified feeding times.

### Animal model

Silica with a diameter of 5 µm (80% of particles) was purchased from Sigma® (S5631). It was incubated at 180 degrees for 16 h to inactivate endotoxin. We used NS to generate a working solution of 50 mg/ml silica suspension before use. Male C57BL/6J mice (6 weeks of age, 22±1 g weight) were anaesthetized with pentobarbital sodium (1%, 50 mg/kg in ddH_2_O) administered by intraperitoneal injection. After the fur on the neck was shaved, the skin was disinfected with 75% alcohol, and an incision (1 cm) was made on the skin along the midline. We bluntly separated the soft tissue in front of the neck to expose the trachea. For the mice in the SiO_2_-7d and SiO_2_-56d groups, we administered 100 µl of silica suspension (50 mg/ml) to the lungs via intratracheal instillation. For the mice in the NS-7d and NS-56d groups, we administered NS via the same method. The mouse skin was sutured and disinfected, and they were closely observed until they recovered. All mice were fasted with water for 6 hours before and after the operation. We observed the mice every 8 hours for the first three days after the operation and once a day afterwards.

### CT scanning

Mice were anaesthetized with inhaled isoflurane (induced concentration 3– 44%, maintained concentration 1–1.5%), and when their breathing was stable, they were placed on the mouse platform, and CT (Hiscan XM Micro CT, Suzhou Hiscan Information Technology Co., Ltd.) detection was performed. The X-ray tube settings were 60 kV and 133 µA, and images were acquired at 50 µm resolution. A 0.5° rotation step through a 360° angular range with 50 ms exposure per step was used. The images were reconstructed with Hiscan Reconstruct software (Version 3.0, Suzhou Hiscan Information Technology Co., Ltd.) and analysed with Hiscan Analyzer software (Version 3.0, Suzhou Hiscan Information Technology Co., Ltd.).

### Pulmonary function test

Mice were anaesthetized by pentobarbital sodium injection as described above. Then, we inserted a tracheal catheter and fastened it to the trachea. After finishing the preparatory work, we tested mouse pulmonary function (Cchord, FEV75 (volume expired in the first 75 ms of fast expiration), IC (inspiratory capacity, volume inspired during slow inspiration), MMEF (mean mid expiratory flow, average flow between 25%-75%), FVC (forced vital capacity, volume expired during fast expiration) and TLC (total lung capacity, FRC+IC) with the Forced Manoeuvres System (EMMS, Hants, UK). Each mouse was tested three times, and the most reliable result (removal of unusually high or unusually low value) was used for analysis. Finally, mice were sacrificed to harvest lung tissue, and the bodies were prepared for cremation by the university.

### Processing of mouse lung tissue for histology

Fresh lung tissues were immediately fixed in 4% paraformaldehyde for 24 h at 4°C. Then, lung tissues were transferred to a 20% sucrose gradient overnight for dehydration. Then, they were transferred to a 30% sucrose gradient overnight for further dehydration. After being fully dehydrated, these lung tissues were stored at -80°C before the next experiment.

### H&E staining

Tissues were cut into 8 µm sections at -20°C by a cryostat (Leica, Germany). The lung tissue sections were stained according to the instructions provided with the H&E staining kit (Biyun Tian, China). Briefly, the tissue sections were washed 3 times in precooled phosphate-buffered saline (1×PBS) and then stained with haematoxylin for 5 minutes and then transferred into acid alcohol fast differentiation solution for 10 seconds. After that, the sections were soaked in tap water for 15 minutes. Next, the cells were stained with Esion for 5 minutes and washed in running water. Then, the sections were dehydrated with 75%, 95%, 100%, 100% alcohol and 100% xylene for 1 min each. Finally, neutral gum and glass slides were used to cover the tissue.

After the sections were dried in air, the images were observed and photographed at 10x and 20x resolution with a microscope (EVOS FL AUTO2, Thermo, America). Images at 10x resolution were captured using EVOS Automated Imaging System software.

### Sirius red staining

All tissue sections were stained according to the instructions of the Sirius Star Chromosome Kit (Abcam, USA). Briefly, 8-µm-thick lung tissue sections were washed in PBS 3 times for 5 minutes and then stained with Sirius red for 1 hour, followed by brief rinsing with 1% acetic acid. Finally, the images were observed and photographed at 10x and 20x resolution with a microscope (EVOS FL AUTO2, Thermo, America). Images at 10x resolution were captured using EVOS Automated Imaging System software.

### ECM processing

#### 1. ECM collection

Fresh lung tissues were cut into 200-µm-thick pieces by cryostat sectioning. After rinsing with PBS, 15 ml of lysis buffer (1% SDS in ddH2O) was added, and the tissue mixture was incubated at room temperature for 1 h on a shaker. Then, the lung tissues were transferred to new lysis buffer and incubated for 1 h again. Next, lung tissues were transferred into another tube with new lysis buffer and incubated overnight at room temperature. On the second day, the sections were incubated with 1% Triton X-100 (diluted with ddH_2_O) at room temperature for 1 h; this incubation was repeated twice. Next, the 1% Triton X-100 solution was renewed and incubated overnight at room temperature. On the third day, the slices were washed with PBS and ddH_2_O in turn for 5 minutes at room temperature. Next, they were put in sodium chloride solution (1 M) for 1 h incubation and then washed with PBS and ddH_2_O in turn for 5 minutes at room temperature. After that, the slices were incubated in solution containing DNase (20 µg/ml) and MgCl_2_ (4.2 mM) at 37°C for 1 h. Finally, the sections were washed with ddH_2_O. Membrane proteins and nuclear proteins were extracted, and lung ECM was obtained.

#### 2. Extraction of ECM proteins

Lung ECM was added to an appropriate amount of SDS lysis buffer (Biyun Tian, China) and incubated at 37°C for 1 h, followed by centrifugation at room temperature at 15000 rcf for 15 min. The supernatant was collected as the ECM protein solution. The ECM protein solution was stored in PBS at 4°C before use.

### Proteomic analysis of the ECM (PXD028194)

#### 1. Experimental procedures

Total protein was extracted from 6 lung ECM samples (3 for the NS-56d group, namely, con111, con116, con117; 3 for the SiO_2_-56d group, namely, M80, M101, M107), and a portion (10 µl) of the protein was used to measure the protein concentration, followed by SDS-PAGE separation. Another part was collected for trypsin digestion and labelled with TMT (Tandem Mass Tags) reagents, con111 with 126, con 116 with 127, con117 with 128; M80 with 129, M101 with 130, M107 with 131. Equal amounts of each labelled sample were mixed, and an appropriate quantity of protein was taken to perform chromatographic separation. Finally, the samples were analysed by LC-MS (liquid chromatography mass spectrometry).

#### 2. Analysis of LC-MS/MS data

The LC-MS/MS raw data was processed using Proteome Discover 2.4 (Thermo, USA). According to the unique peptide ≥ 1, keep any group of samples with the expression value ≥ 50% of the protein. Then, the missing values were imputed with the mean of the protein expression in corresponding group. Next, the data was median normalized and log2-transformed to obtain credible proteins. Then we performed statistical and visual display of these proteins using R software (version 4.2) ggplot2 package (version 3.2.2), including principal component analysis (PCA), sample correlation analysis (Extended Data Fig. 2a), sample hierarchical cluster analysis (Extended Data Fig. 2b), visual display of data after standardization (Extended Data Fig. 2c) and density plot (Extended Data Fig. 2d).

Based on credible proteins, we performed Student’s t test to identify significant difference of proteins in NS-56d group and SiO_2_-56d group. Fold change (FC) is used to evaluate the expression level of a certain protein between samples. The p-value (P) calculated by the t-test shows the significance of the difference between samples. Difference screening conditions are FC≥2.0 and P≤0.05. The clustering heat map based on R software (version 4.2) pheatmap package (version 1.0.12) can be used for quality control of standardized experimental data and data display after enrichment of differential data. Generally, samples of the same group can appear in the same cluster through clustering.

For the identified proteins, extract the annotation information based on such as Uniprot databases. After obtaining the differentially expressed proteins (FC≥2, P≤0.05), the GO and KEGG functional enrichment analyses of up-regulated proteins were performed with R software (version 4.2) ggplot2 package (version 3.2.2).

### Western blotting

The proteins were treated with loading buffer at 100°C for 5 min. Then, equal amounts of proteins (20 µg) of lung ECM were added to a 12% SDS-PAGE gel, and after separation, the proteins were transferred to a 0.25 µm PVDF membrane. Next, the PVDF membrane was placed in 5% skim milk at room temperature for one hour to block nonspecific binding sites. Then, the membranes were incubated in primary antibody (anti-pro/mature surfactant protein B, ab40876, Abcam, 1:5000) at 4°C overnight. After being washed 4 times with TBST at room temperature for 8 min, the PVDF membranes were incubated with HRP-conjugated goat anti-mouse or anti-rabbit secondary antibodies (according to what primary antibody origination was used) for 1 h at room temperature. For the detection of IgA, we used goat anti-mouse IgA-HRP (Southern Biotech, SBA-1040-05, 1:5000) as both the primary antibody and secondary antibody. After unbound secondary antibodies were fully washed away with TBST, the signal was detected with ECL chemiluminescence solution (Millipore, America), and photos were taken with a chemiluminescence detection system (Tanon Scientific & Technology Co., China). For lung ECM samples, β-actin was used as an internal reference protein. For MLE-12 cell line samples, GAPDH was used as an internal reference protein.

### Spatial transcriptomics(GSE183683)

#### 1. Sample collection

For the model group, mice with obvious fibrotic lesions on CT were thought to be a successful model. Lung tissues were trimmed near the hilum in the horizontal direction and then immediately frozen in OCT on dry ice. These samples were stored at -80°C before next step.

#### 2. Spatial transcriptomics sequencing

##### 2.1 Staining and imaging

Cryosections were cut at 10 μm thickness with a cryostat (Leica, Germany) and mounted onto GEX arrays. The arrays were placed on a Thermocycler Adaptor with the active surface facing up and then incubated for 1 min at 37°C, fixed for 30 min with methyl alcohol at -20°C, and stained with H&E. Bright field images of the whole slide were acquired on a Leica DMI8 whole-slide scanner at 10x resolution.

##### 2.2 Permeabilization and reverse transcription

Spatial gene expression analysis was performed using the Visium Spatial Gene Expression Slide and Reagent Kit (10x Genomics, PN-1000184). For each well, a slide cassette was used to create leakproof wells for adding reagents. Then, 70 μl of permeabilization enzyme was added, and the samples were incubated at 37°C. For the NS-7d, SiO2-7d and NS-56d samples, the incubation time was 24 min. However, for the SiO2-56d group, because of severe lung fibrosis, a 30-min incubation time was used. Each well was washed with 100 μl of SSC, and 75 μl of RT master mix was added for cDNA synthesis (65°C, 15 minutes, hold at 4°C).

##### 2.3 cDNA library preparation for sequencing

At the end of first-strand synthesis, RT Master Mix was removed from the wells.

Then, 75 μl of 0.08 M KOH was added and incubated for 5 min at room temperature. Then, KOH was removed from the wells, which were washed with 100 µl of EB buffer. Then, 75 μl of Second Strand Mix was added to each well for second-strand synthesis. cDNA amplification (98°C 3 min; 98°C 15 s, 63°C 20 s, 72°C 1 min, 14 cycles; 72°C 1 min, hold at 4°C) was performed on a S1000TM Touch Thermal Cycler (Bio-Rad). According to the manufacturer’s instructions, Visium spatial libraries were constructed using a Visium spatial library construction kit (10x Genomics, PN-1000184). The libraries were sequenced using an Illumina NovaSeq6000 sequencer with a sequencing depth of at least 100,000 reads per spot with a paired-end 150 bp (PE150) reading strategy (performed by CapitalBio Technology, Beijing).

#### 3. Analysis of spatial transcriptomics data

The 10X Space Ranger software, which process, align and summarize unique molecular identifier (UMI) counts against mmu10 mouse reference genome for each spot, was used to generate feature-barcode matrix. Only spots overlaying tissue sections were retained for further analysis. Unsupervised clustering was based on graph based algorithm with 10 principal components. The *t*-distributed stochastic neighborhood embedding (*t*-SNE) was performed to visualize the spots in two-dimensional space. Spatial feature expression plots were generated by Loupe Browser 4.1.0.

### Single-cell sequencing (GSE183682)

#### 1. Sample collection

For the model group (mice instilled with SiO2 suspension), mice with obviously high-density shadows on CT were included. Lung samples for single-cell sequencing were collected from four mice, namely, the NS-7d, SiO_2_-7d, NS-56d, and SiO_2_-56d samples. Whole lungs of each mouse were removed within 2 min of euthanasia and quickly washed in precooled PBS 3 times.

#### 2. scRNA-seq

##### 2.1 Cell capture and cDNA synthesis

Whole lung tissue was cut into small pieces (approximately 1 mm) and dissociated into single cells using Lung Dissociation Kit (Miltenyi Biotech, 130-095-927, Germany). With the Single-Cell 5’ Library and Gel Bead Kit (10x Genomics, 1000169) and Chromium Single-Cell G Chip Kit (10x Genomics, 1000120), a cell suspension (300–600 living cells per microlitre determined by CountStar) was loaded onto a Chromium single-cell controller (10x Genomics) to generate single-cell gel beads in emulsion (GEMs) according to the manufacturer’s protocol. In short, single cells were suspended in PBS containing 0.04% BSA. Approximately 20,000 cells were added to each channel, and the target cell recovery was estimated to be approximately 10,000 cells. Captured cells were lysed, and the released RNA was barcoded through reverse transcription in individual GEMs. Reverse transcription was performed on a S1000TM Touch Thermal Cycler (Bio-Rad) at 53°C for 45 min, followed by 85°C for 5 min and a hold at 4°C. cDNA was generated and then amplified, and quality was assessed using an Agilent 4200 system (performed by CapitalBio Technology, Beijing).

##### 2.2 scRNA-seq library preparation

The scRNA-seq libraries were constructed using the Single-Cell 5’ Library and Gel Bead Kit, Single Cell V(D)J Enrichment Kit, Human T Cell (1000005) and Single Cell V(D)J Enrichment Kit according to the manufacturers’ instructions. The libraries were sequenced using an Illumina NovaSeq6000 sequencer with a sequencing depth of at least 100,000 reads per cell with a paired-end 150 bp (PE150) reading strategy (performed by CapitalBio Technology, Beijing).

#### 3. Data preprocessing

##### 3.1 Analysis of scRNA-seq data

Cell barcode filtering, alignment of reads and UMI counting were performed with Cell Ranger 4.0.0 (https://www.10xgenomics.com/). The scRNA-seq data for four samples was combined though Cellranger aggr. Principal component analysis (PCA) is performed on the normalized data. Unsupervised clustering was performed by Cellranger reanalyze using graph based algorithm. The top 10 principal components was used for clustering and *t*-SNE projections. The differential expression of genes between clusters were computed by sSEq and edgeR based method. The GO and KEGG functional enrichment analyses of marker genes (FC≥2, P≤0.05) in cluster 1 and 6 were performed by R package Metascape.

##### 3.2 Cell type annotation

Cell types were determined by clustering and marker gene expression. Cluster 1, 3 and 15 highly expressed genes *Sftpc*, *Sftpb* and *S*ftpa1, which were defined as AT2-like cell. Cluster 6 highly expressed genes *Itgax*, *Csf1r* and *Ly86*, and was inferred to be macrophage. Monocyte was made up two clusters (cluster 4 and 8), which highly expressed *Ccr2* and *Csf1r*. Other clusters highly expressed markers specific for dentritic cell (*Ear1*, *Plet1*), clara cell (*Sftpb*, *Sftpc*, *Scgb3a1*), AT1 cell (*Cldn3*, *Epcam*), Ccl3- Ccl4- neutrophils (*Cxcr2*, *Stfa2l1*), Ccl3+ Ccl4+ neutrophils (*Il1f9*, *Ccl3*, *Ccl4*), T cell (*Thy1*, *Cd8a*), B cell (*Cd79a*, *Cd79b*, *Cd19*), endothelial cell (*Cdh5*, *Pecam1*, *Clec14a*), fibroblast (*Col1a1*, *Col3a1*, *Col6a2*), red blood cell (*Hba-a1*, *Hba-a2*)

#### 4. BCR and TCR sequence reconstruction

Using a Chromium Single-Cell V(D)J Enrichment kit, we reconstructed full-length TCR/BCR V(D)J segments from amplified cDNA in 5′ libraries via PCR amplification of sc-RNA seq following the manufacturer’s protocol (10x Genomics). Using the Cell Ranger (v.3.0.2) vdj pipeline coupled with the mouse reference genome mm10, we conducted demultiplexing, gene quantification and TCR/BCR clonotype assignment. Cells with at least one complete TCR α-chain (TRA) or TCR β-chain (TRB) were retained for TCR analysis, and only cells with at least one complete heavy chain (IGH) or one light chain (IGK or IGL) were retained for BCR analysis. Each unique TRA(s)-TRB(s) pair or IGH(s)-IGK/IGL(s) pair was defined as an effective clonotype for further clonal expansion analysis. Cells with the same clonotype were considered clonal cells. Based on barcode information, cells with a TCR or BCR clonotype were projected on UMAP plots.

### RNAscope

#### 1. Preparation of lung tissue section

Lung tissues harvested from mouse of NS-56d and SiO_2_-56d (mouse and modeling methods are the same as in scRNA sequencing) were immediately frozen in OCT on dry ice. These samples were stored at -80°C before next step.

#### 2. RNAscope in situ hybridization

All RNAscope experiments are performed strictly in accordance with ACD’s instructions. After being equilibrated at -20°C for 1h, the tissues were cut into 12 µm sections at -20°C by a cryostat (Leica, Germany) and mounted onto SuperFrost Plus slides (Fisher, Scientific, 12-550-15). The tissue sections were air-dried at -20°C for 20 minutes and then immediately transferred into 4% paraformaldehyde (precooled at 4°C ) for 15 minutes. Next, the tissue sections were dehydrated in 50% ethanol, 70% ethanol, 100% ethanol and 100% ethanol in order at room temperature, each for 5 minutes. The sections were air-dried for 5 minutes at room temperature and then the boundaries of sections were drawn on slides using an ImmEdge hydrophobic pen. After the hydrophobic boundaries were dried, tissue sections were incubated with hydrogen peroxide solution for 10 minutes at room temperature, followed by briefly washed twice in PBS at room temperature. Tissue sections then were incubated with protease IV for 15 minutes at room temperature followed by briefly washed twice in PBS at room temperature.

The tissue sections were placed in humidity box and incubated with a mixture of *Sftpc* probe (C3, 570) and *Igha* probe (C2, 520) ( 1:50 diluted with probe diluent) at 4°C for 2 hours, followed by washed twice in 1×washing buffer at room temperature for 2 minutes. Then tissue sections were incubated with AMP-1, AMP-2, AMP-3 reagents for 30, 30, 15 minutes respectively at 40°C. After each amplification, the sections were washed twice in 1×washing buffer at room temperature for 2 minutes. The sections were incubated with HRP-C2 reagent at 40°C for 15 minutes, followed by washed twice in 1×washing buffer at room temperature for 2 minutes. After that, the sections were incubated with Opal 520 (1:1000 diluted with TSA diluent) at 40°C for 30 minutes, followed by washed twice in 1×washing buffer at room temperature for 2 minutes. The sections were incubated with HRP-C3 reagent at 40°C for 15 minutes, followed by washed twice in 1×washing buffer at room temperature for 2 minutes. After that, the sections were incubated with Opal 570 (1:1000 diluted with TSA diluent) at 40°C for 15 minutes, followed by washed twice in 1×washing buffer at room temperature for 2 minutes.

Finally, the sections were incubated with DAPI reagent for 30s. After briefly washed in PBS, the sections were covered with Prolong Gold Antifade mounting medium (Thermo Fisher Scientific, P36930, USA). All the slides were fully air-dried and then stored in dark at 4°C.

#### 3. Image capture

Images were captured using a confocal (Olympus FV1000, Japan). We observe the overall condition of the sample through a 10× objective lens. The detailed pictures were taken with a 60× objective oil lens. Images of different channels were taken separately and merged using Olympus Fluoview software.

### Cell culture

MLE12 mouse lung epithelial cells (ScienCell) were cultured in DMEM (Gibco) supplemented with 10% FBS (Corning) and 1% penicillin-streptomycin (Gibco) in a 5% CO2 and 37°C incubator (Thermo).

### Protein extraction

Cells were washed 3 times with precooled PBS and then lysed with RIPA lysis buffer (Beyotime Biotechnology, China) supplemented with a protease inhibitor cocktail (Beyotime Biotechnology, China) on ice for 1 h, followed by centrifugation at 12000 rpm for 15 min. Then, the protein concentration in the supernatant was quantified using a BCA assay (Beyotime Biotechnology, China), and the extracted protein was heated for 5 min at 100°C. The protein solution was stored at -80°C before use.

### scRNA-seq and spatial RNA-seq integration

In order to obtain the composition of cell types in each spot, we used anchor-based integration method in Seurat v3.2, which could transfer the label of scRNA-seq reference to a spatial RNA-seq query. First, both the scRNA-seq and spatial RNA-seq data were normalized with the SCTransform function. Then, the cell type prediction probabilities score were calculated for each spot through FindTransferAnchors and TransferData functions. Finally, the spot is assigned to the cell type with the highest score.

---

**Extended Data Fig. 1:**
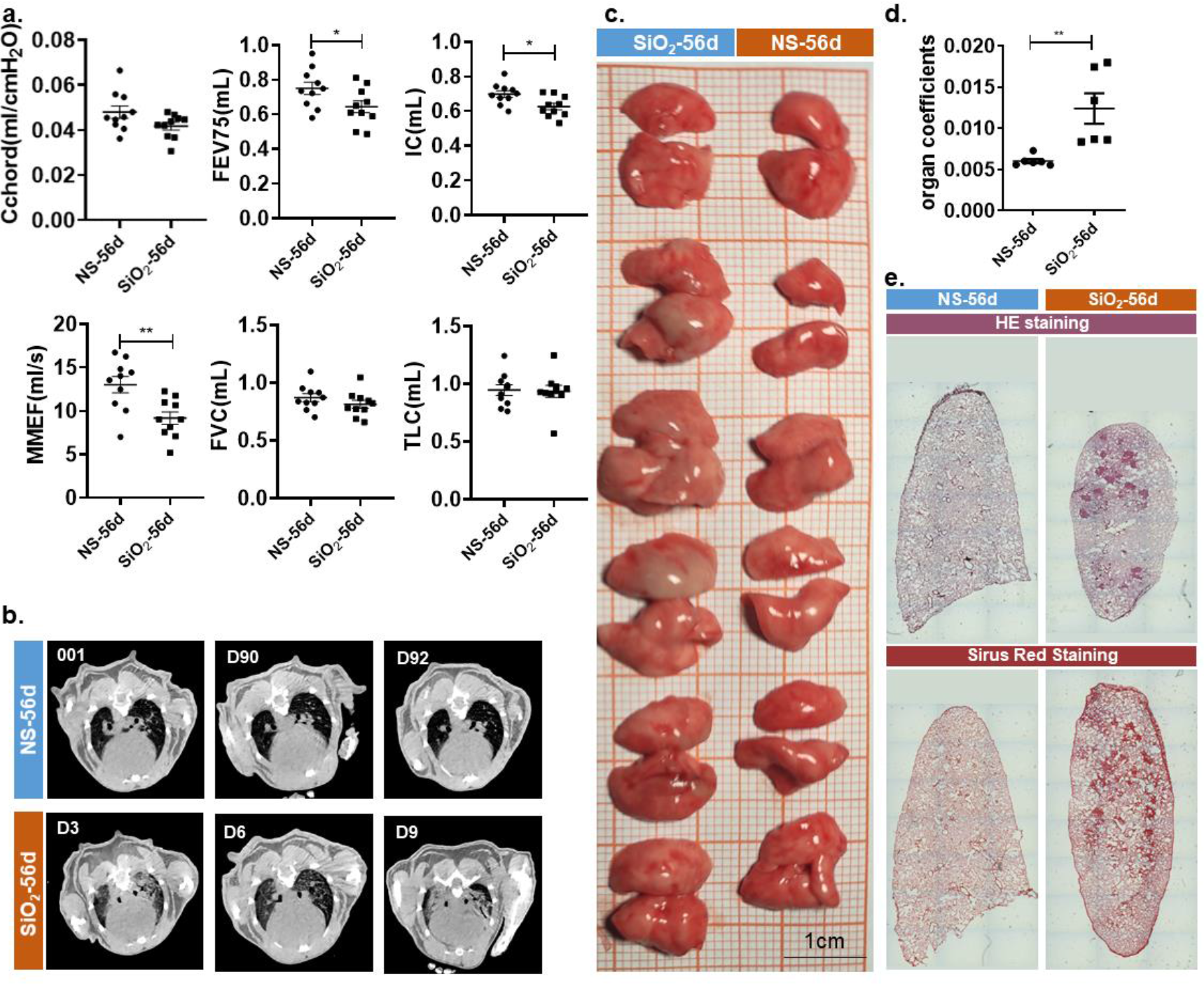
Validation of fibrosis stage. a, Results of lung function on day 56 after instillation. Chord (ml/cmH2O), chord compliance; FEV75 (ml), the volume expired in the first 75 ms of fast expiration; IC (ml), inspiratory capacity, the volume inspired during slow inspiration; MMEF (ml/s), mean midexpiratory flow, the average flow between 25%-75% FVC; FVC (ml), forced vital capacity, the volume expired during fast expiration; TLC (ml), total lung capacity, FRC+IC. b, Chest CT images of NS-56d and SiO_2_-56d. c, Photos of fresh lung. Above is NS-56d; below is SiO_2_-56d. d, Organ coefficients of lung. e, H&E staining and Sirius red staining of NS-56d and SiO_2_-56d.

**Extended Data Fig. 2:**
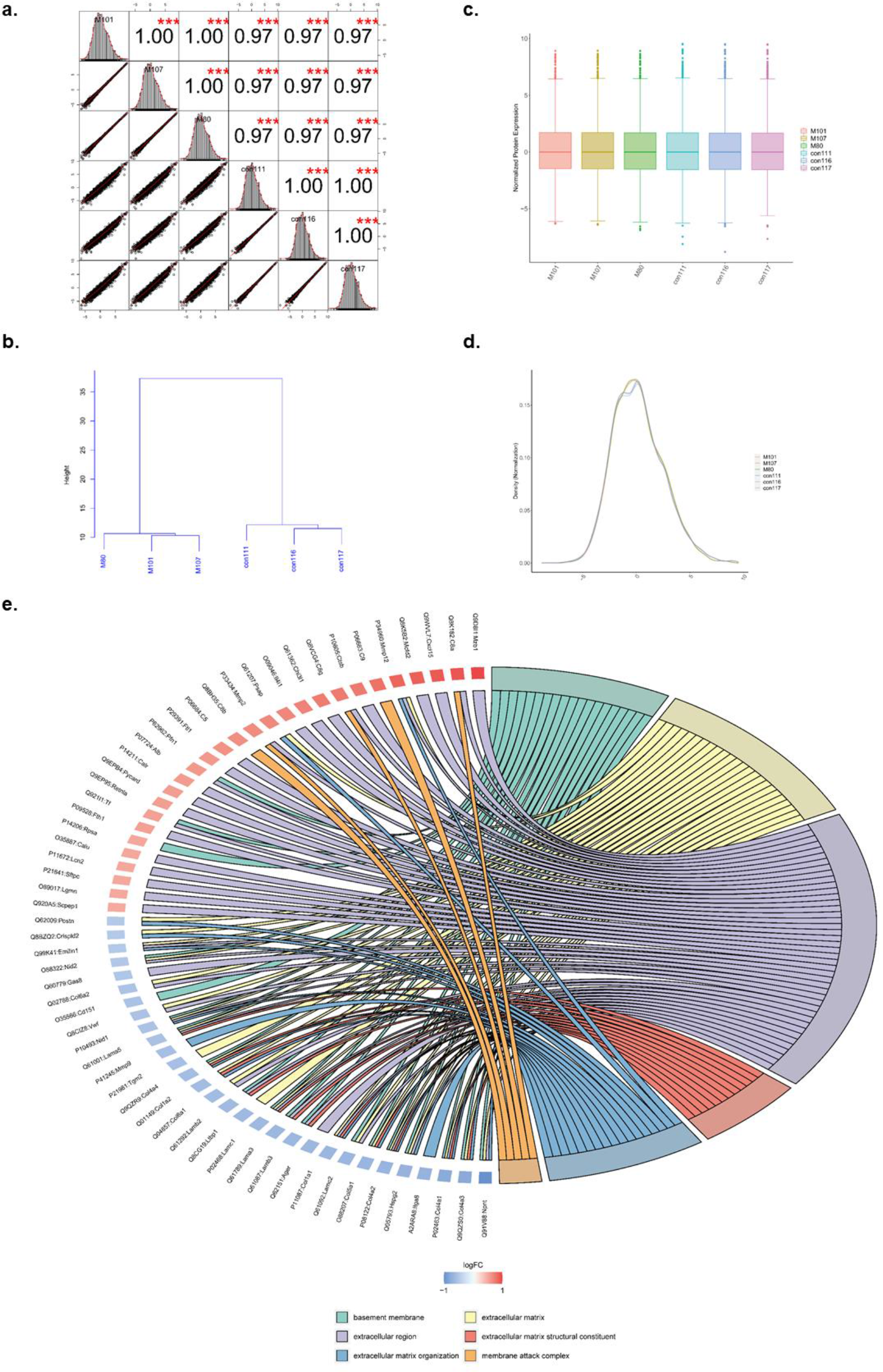
Qualitative-quantitative analysis of proteins. a, Correlation analysis of identified proteins in 6 samples. The degrees of correlation reflect groups and repeatability of samples. The numbers (top right) indicate the correlation of two samples. * indicates significance (* p < 0.05, ** p < 0.01, *** p < 0.001). Scatter plots show the expression of proteins in two samples (bottom left). The red curve is the fitting curve. The slope indicates the correlation between two samples. Diagrams on the diagonal show the distribution of protein expression in one sample. M80, M101 and M107 are samples of SiO2-56d; con111, con116 and con117 are samples of NS-56d. b, Hierarchical clustering dendrogram of sample Euclidean distance indicating similarity of samples. c, Boxplots show the degree of concentration for each data point. The flatter the box is, the more concentrated the data are. The shorter the dotted line is, the more concentrated the data are. d, Standardized density diagram. The data were close to a normal distribution after standardization. X axis, the value of the sample expression; Y axis, the probability density. e, Chordal graph shows the relationship between GO terms (right) and DEGs (left).

**Extended Data Fig. 3:**
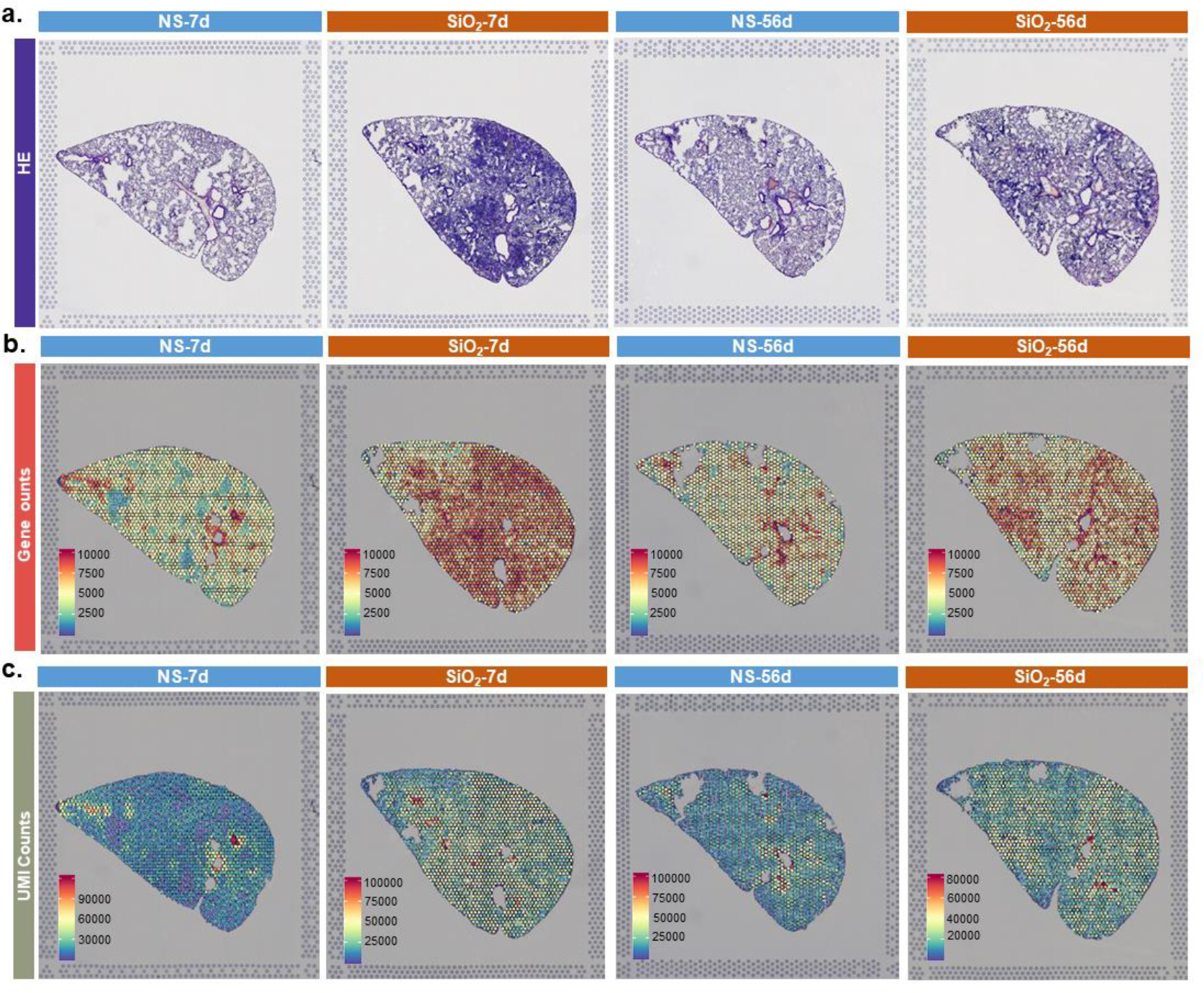
Quality of special transcriptome data. a, H&E staining of lung sections. b, UMI counts displayed on tissue sections. c, Gene counts displayed on tissue sections.

**Extended Data Fig. 4:**
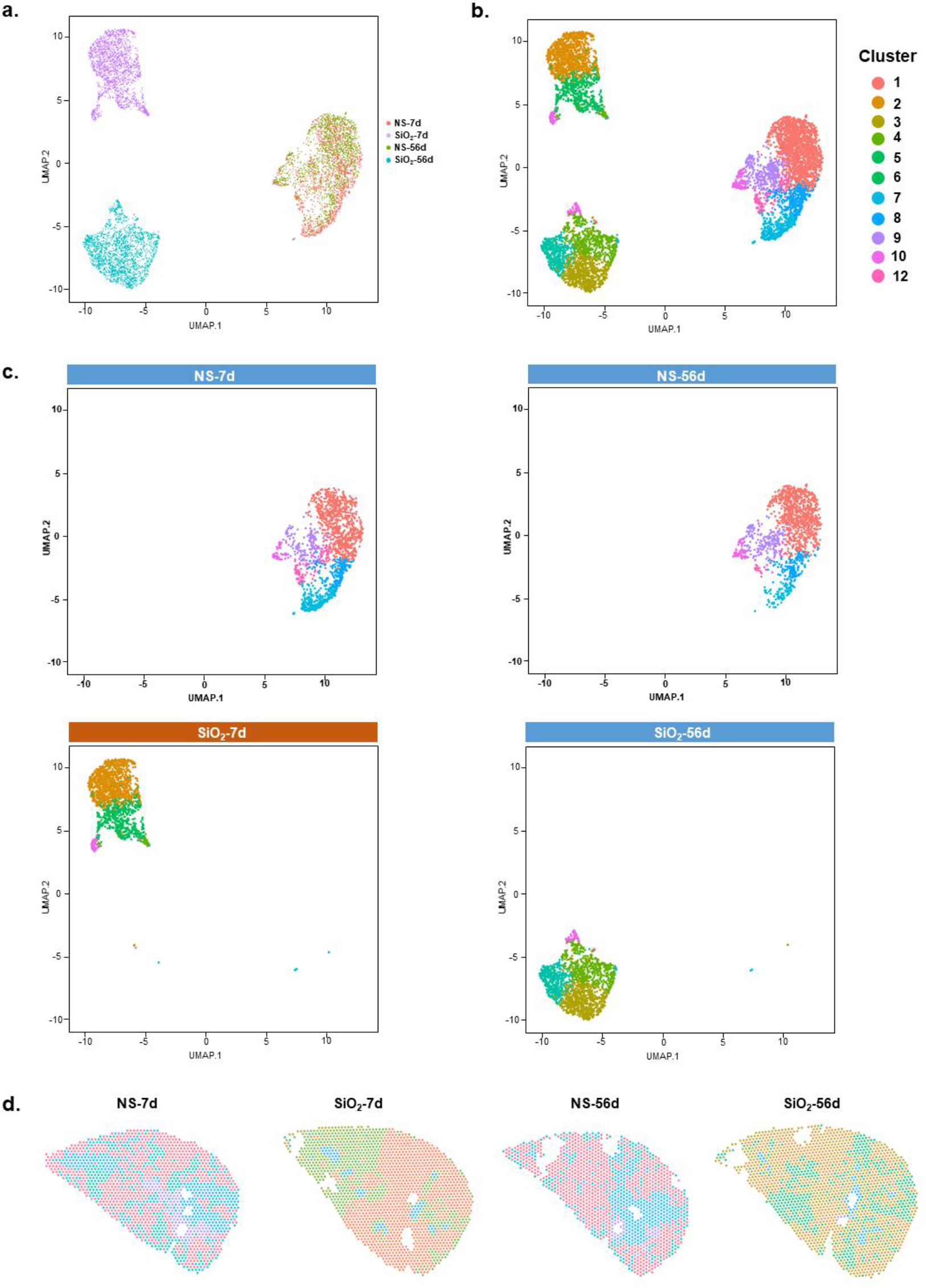
Unbiased clustering of spatial transcriptome spots in four samples. a, UMAP graph of spots coloured by four samples. b, Clusters of spatial transcriptome spots on UMAP: 11 clusters in total, labelled in different colours. c. UMAP graph of individual sample, separated from (b). d, Clusters of spatial transcriptome spots shown in the tissue space.

**Extended Data Fig. 5:**
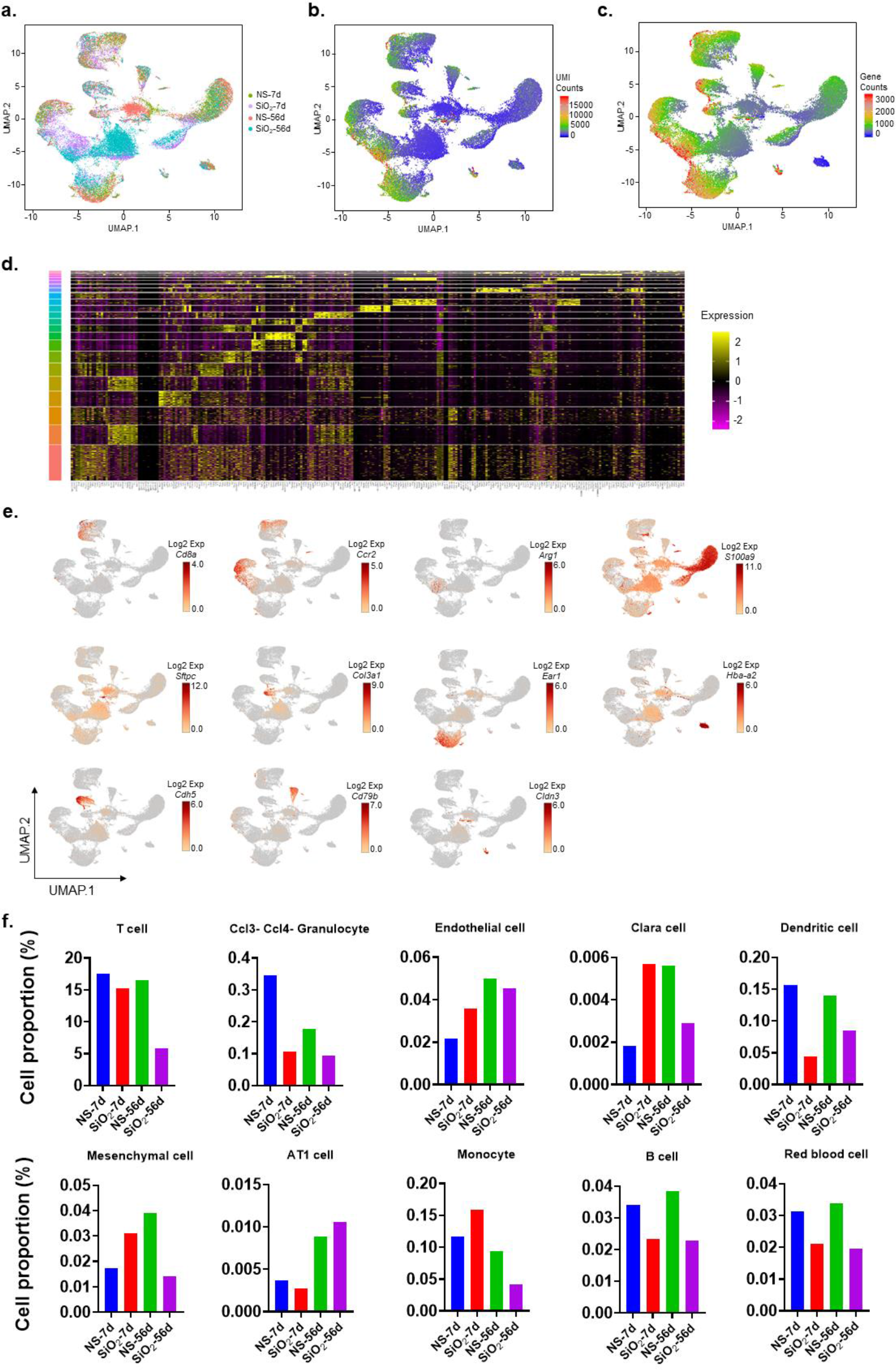
Quality of scRNA-seq data and cell proportions of different cell types. a, UMAP projections of all cells from four samples. Samples are labelled in different colours. b, UMAP graph of UMI counts. c, UMAP graph of gene counts. d, Heat map of the DEGs among the 24 cell clusters. DEGs, differentially expressed genes. AT2-1, AT2-2, AT2-3 and B cells are genes of particular interest. e, UMAP graph of canonical cell markers used to identify 15 cell types. f, Cell proportions of different cell types in four samples.

**Extended Data Fig. 6:**
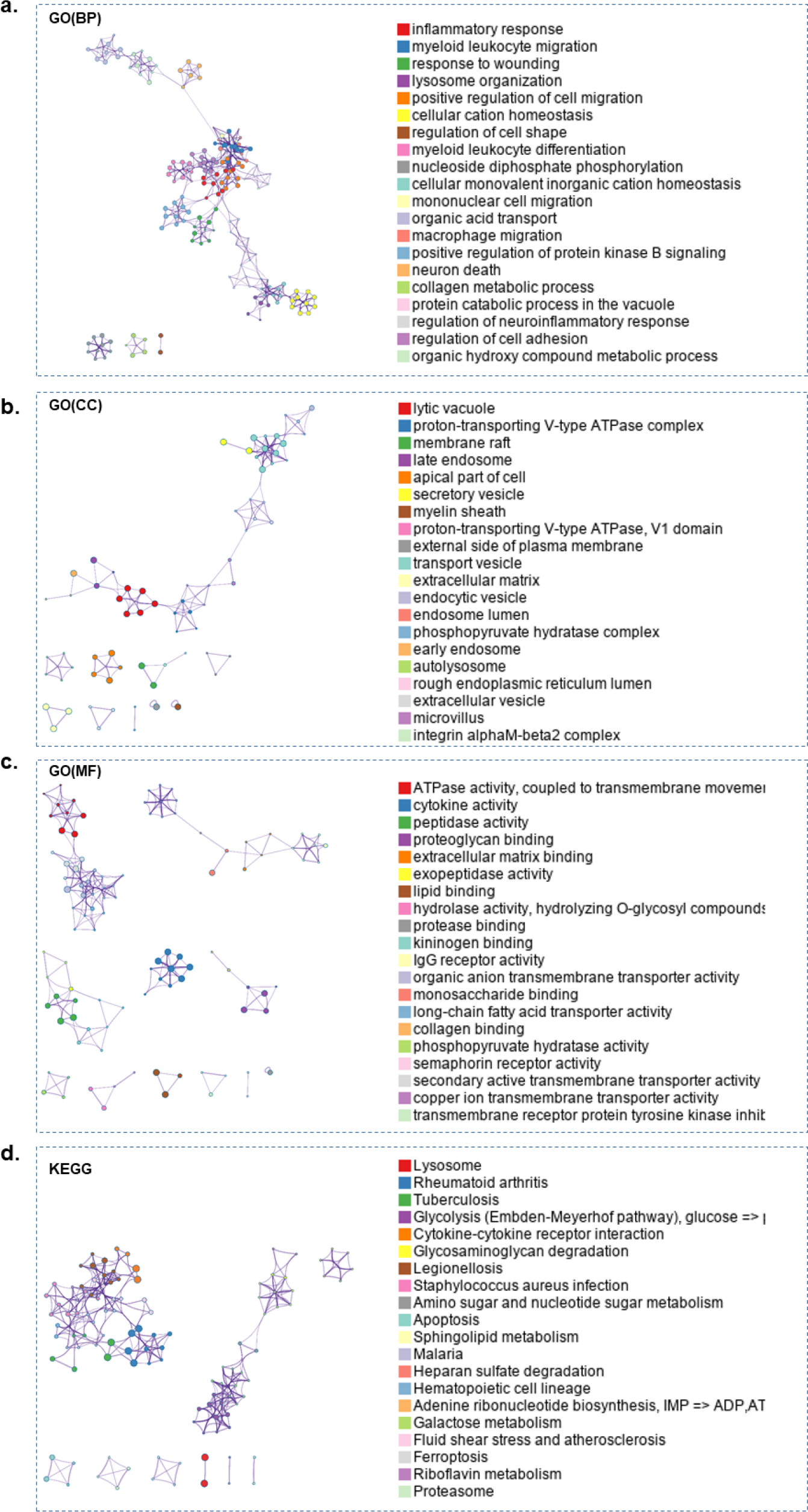
Enrichment analysis of DEGs in macrophages. a-c , GO enrichment of DEGs in macrophages. d, KEGG enrichment of DEGs in macrophages.

**Extended Data Fig. 7:**
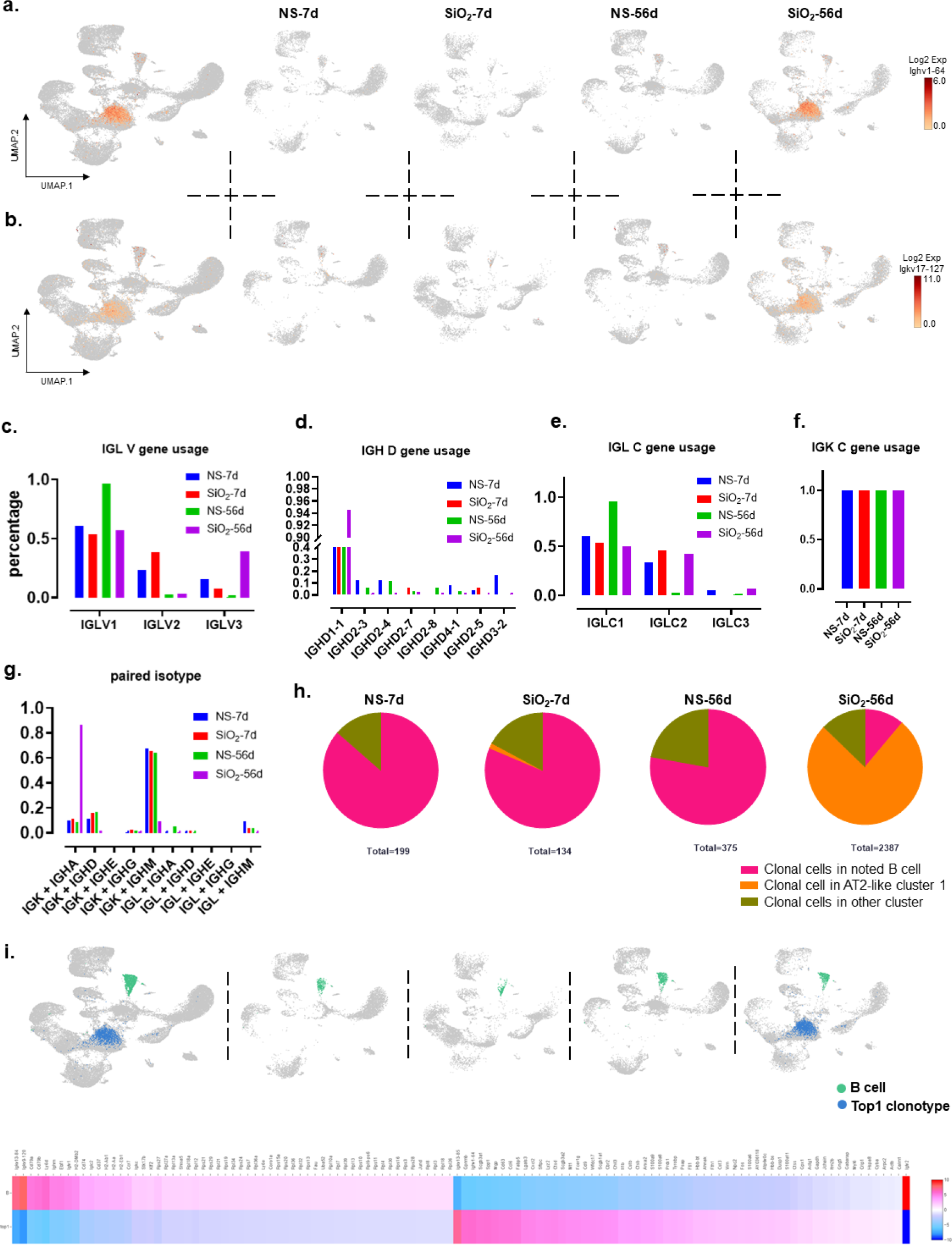
Clonal expansion in SiO_2_-56d. a, Expression of Ighv1-64 in the UMAP graph. b, Expression of Igkv17-127 in the UMAP graph. c, Usage of IGLV genes. d, Usage of IGHD genes. e, Usage of IGLC genes. f, Usage of IGKC genes. g, Paired isotypes in four samples. h, Clonal cells in AT2-like cluster 1 occupied approximately 75% of all SiO_2_-56d. i, B cells and top1 clonal cells. displayed in the UMAP graph. Heat map shows the DEGs between B cells and top1 clonal cells.

**Extended Data Fig. 8:**
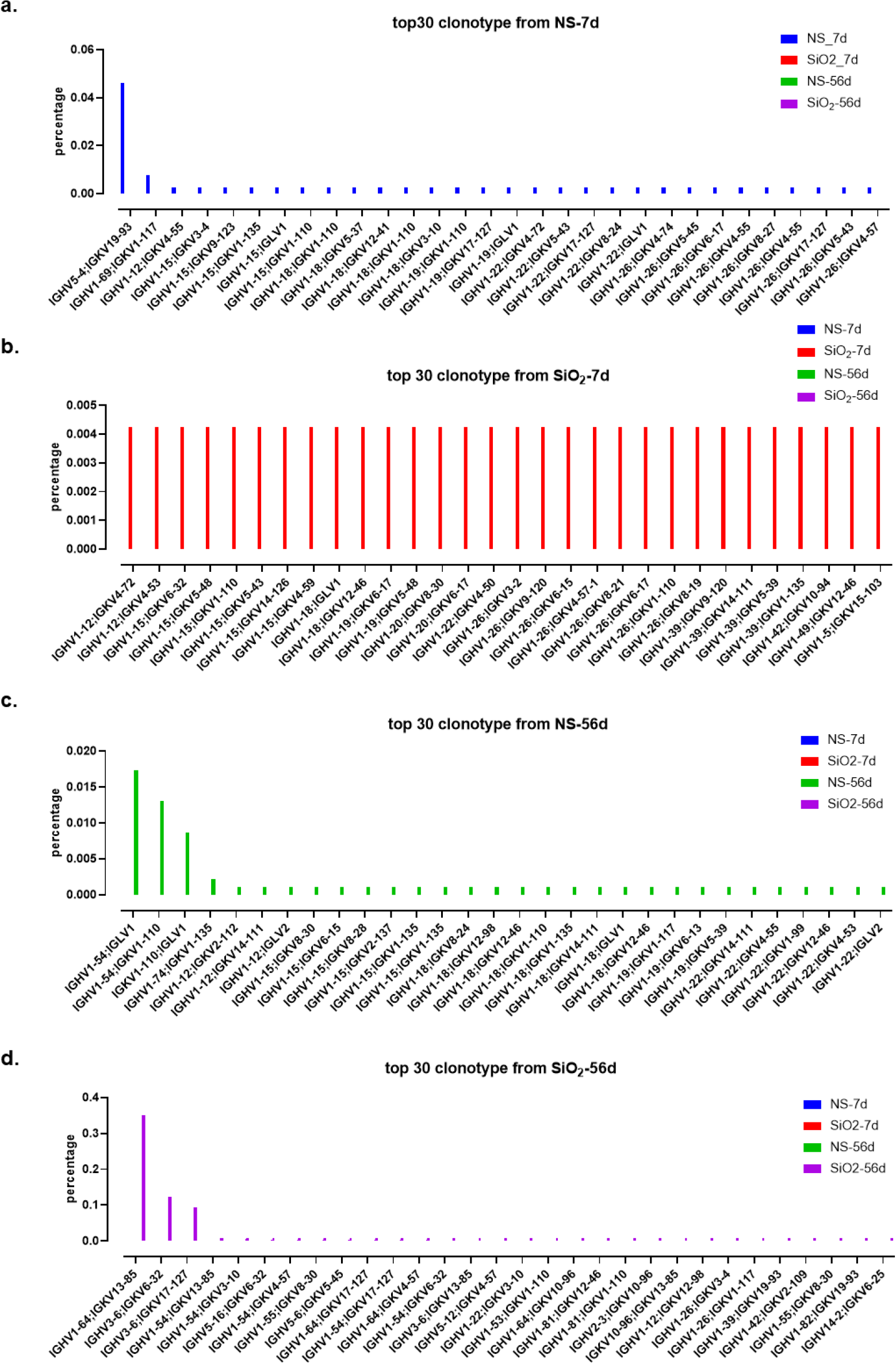
Top 30 clonotypes in four samples. a-d , Each statistical profile shows the percentage of the top 30 clonotypes from one sample, and the percentages of these clonotypes in the other three samples are shown at the same time. Clonotypes in (a) NS-7d; clonotypes in (b) SiO_2_-7d; clonotypes in (c) NS-56d; clonotypes in (d) SiO_2_-56d.

**Extended Data Fig. 9:**
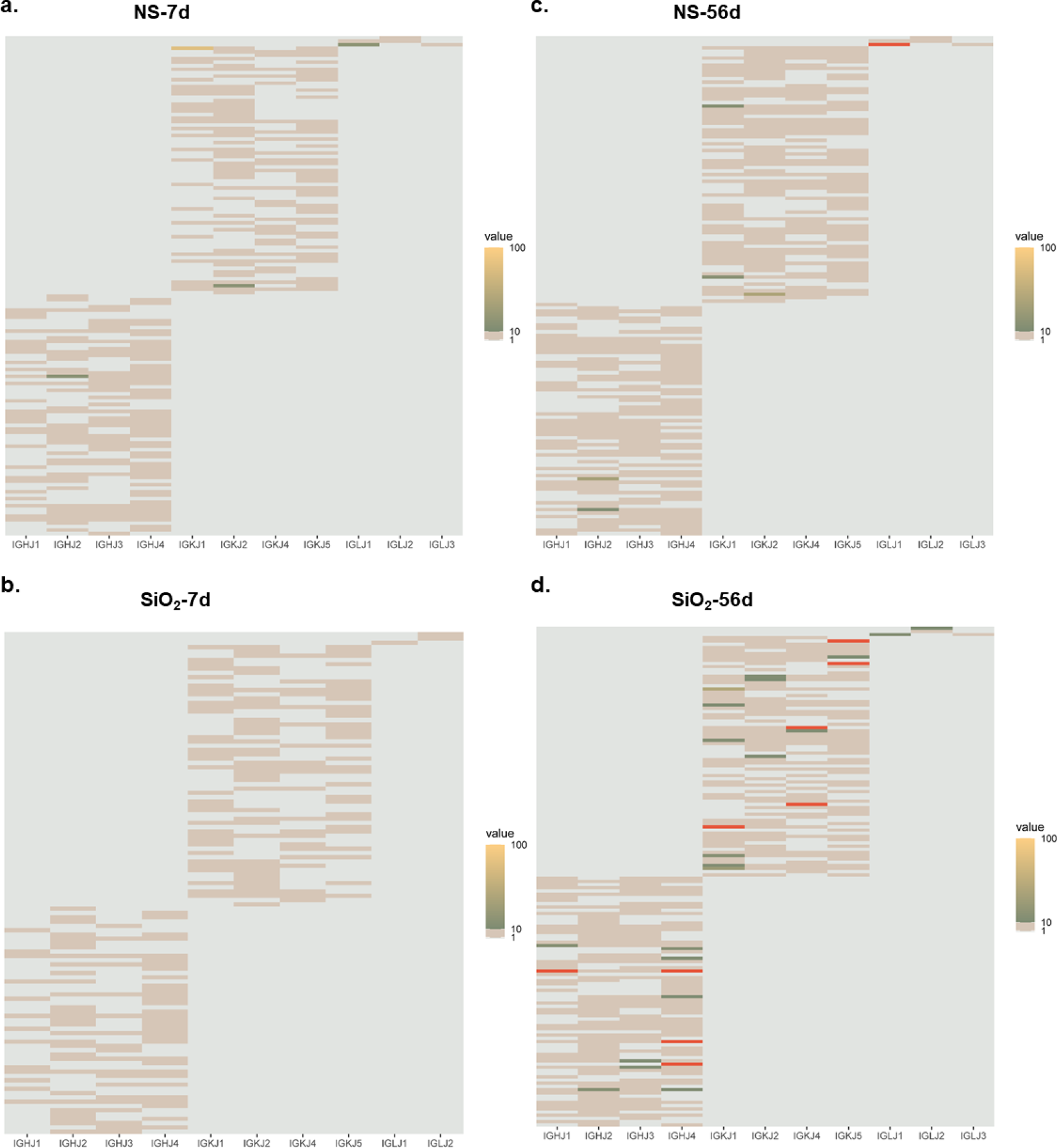
The frequency of V-J gene pairs in four samples. a-d, Heat map indicating the difference in V-J gene rearrangement across four samples. (a) for NS-7d; (b) for SiO_2_-7d; (c) for NS-56d; (d) for SiO_2_-56d.

**Extended Data Fig. 10:**
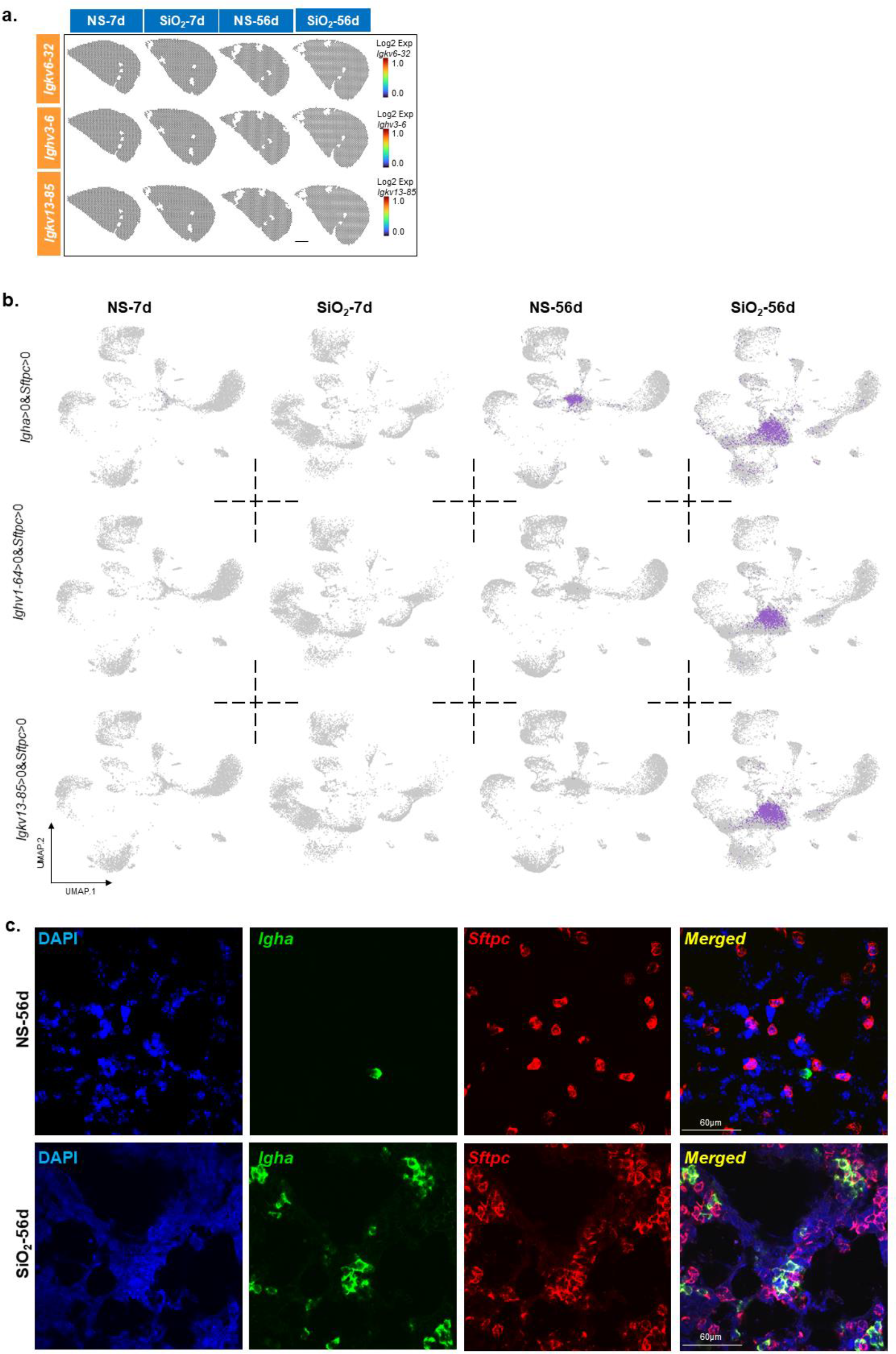
V gene preference with no general sense. a, Expression of Ighv1-64 and Igkv13-85 in the spatial transcriptome of four samples. b, Igha (top), Ighv1-64 (middle) and Igkv13-85 (bottom) coexpressed separately with Sftpc. c, Detailed picture in red frame of Fig. 4h. DAPI, blue; Sftpc, red; Igha, green.

**Extended Data Fig. 11:**
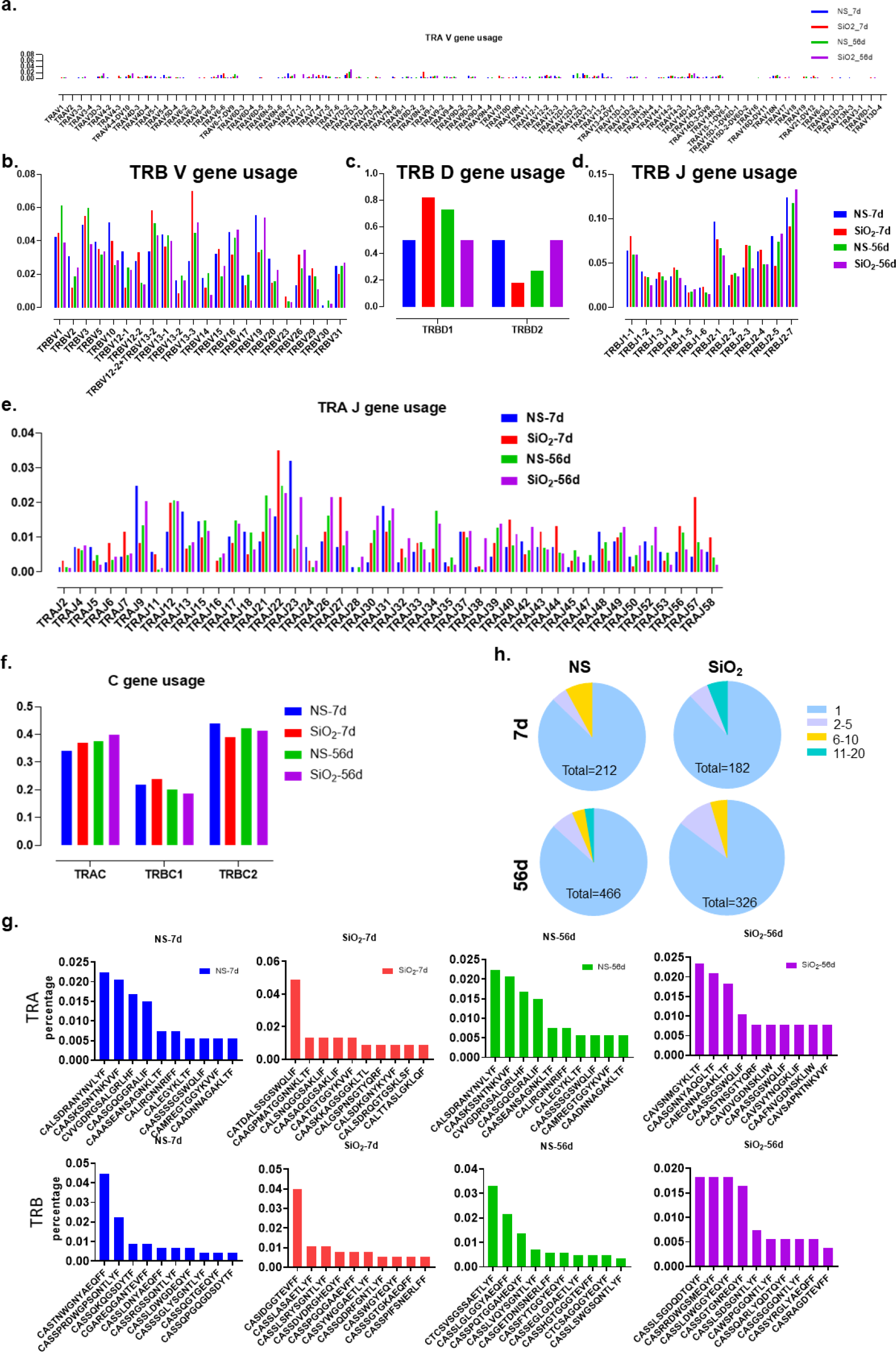
The usage of the V(D)J gene and CDR3 sequence of TCR. a, Usage of TRAV genes. b, Usage of TRBV genes. c, Usage of TRBD genes. d, Usage of TRBJ genes. e, Usage of TRAJ genes. f, Usage of TRAC and TRBC genes. g, Top 10 CDR3 usage in four samples. h, Clonal expansion in four samples.

**Extended Data Fig. 12:**
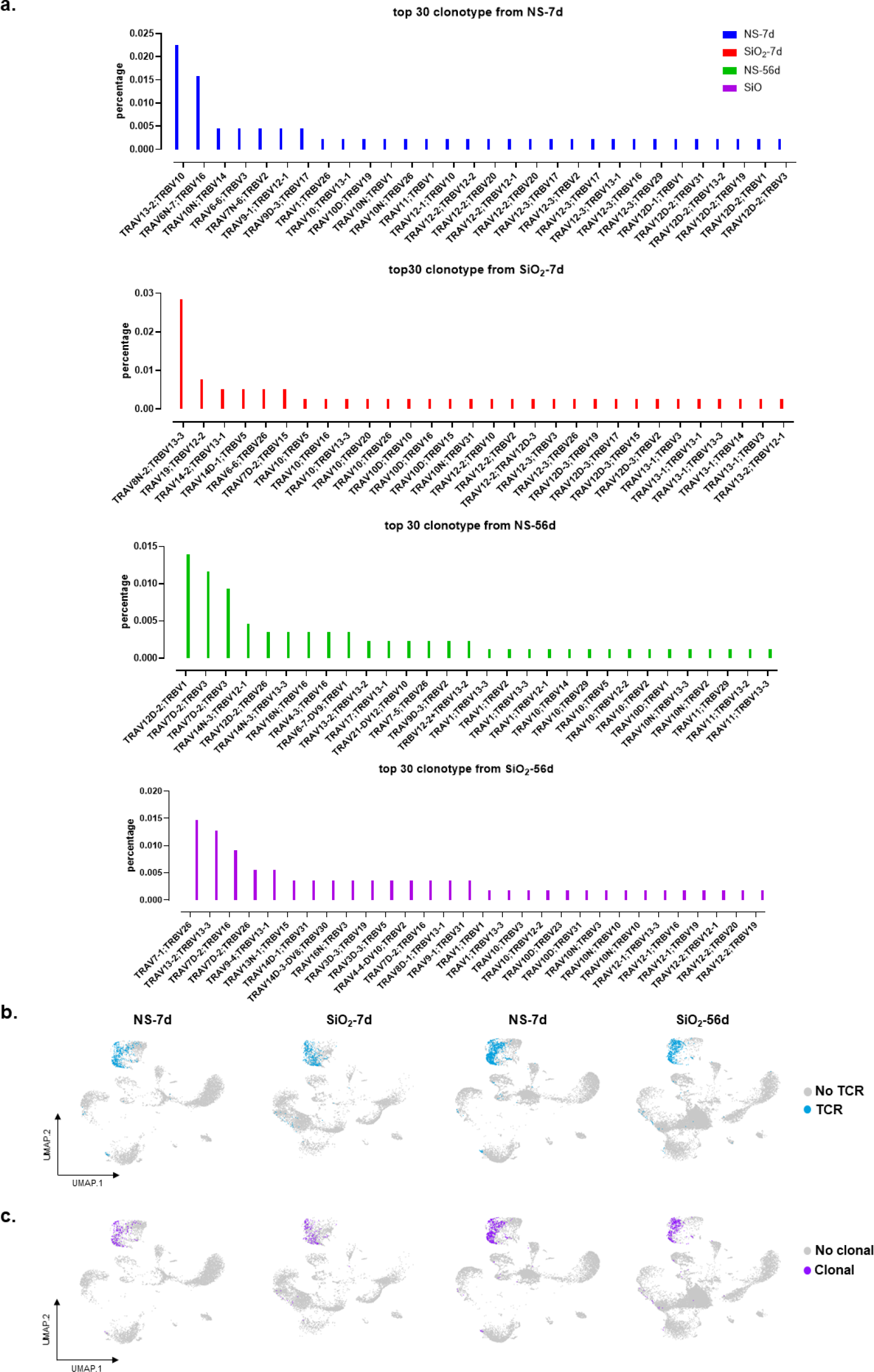
TCR clonal expansions in four samples. a, Top 30 clonotypes in each samples. Every statistical profile shows the percentage of the top 30 clonotypes in one sample and the percentages of these clonotypes in the other three samples. b, UMAP graph of cells with TCRs. c, UMAP graph of cells with clonotypes.

## Notes

### Competing Interest Statement

The authors have declared no competing interest.

## References

Asp, M., Giacomello, S., Larsson, L., Wu, C.L., Furth, D., Qian, X.Y., Wardell, E., Custodio, J., Reimegard, J., Salmen, F., et al. (2019). A Spatiotemporal Organ-Wide Gene Expression and Cell Atlas of the Developing Human Heart. Cell 179, 1647-+. 10.1016/j.cell.2019.11.025.

Babbage, G., Ottensmeier, C.H., Blaydes, J., Stevenson, F.K., and Sahota, S.S. (2006). Immunoglobulin Heavy Chain Locus Events and Expression of Activation-Induced Cytidine Deaminase in Epithelial Breast Cancer Cell Lines. Cancer research 66, 3996–4000. 10.1158/0008-5472.can-05-3704.

Barkauskas, C.E., Cronce, M.J., Rackley, C.R., Bowie, E.J., Keene, D.R., Stripp, B.R., Randell, S.H., Noble, P.W., and Hogan, B.L.M. (2013). Type 2 alveolar cells are stem cells in adult lung. Journal of Clinical Investigation 123, 3025–3036. 10.1172/JCI68782.

Ferreira, R.M., Sabo, A.R., Winfree, S., Collins, K.S., Janosevic, D., Gulbronson, C.J., Cheng, Y.H., Casbon, L., Barwinska, D., Ferkowicz, M.J., et al. (2021). Integration of spatial and single-cell transcriptomics localizes epithelial cell-immune cross-talk in kidney injury. JCI insight 6. ARTN e147703 10.1172/jci.insight.147703.

Han, X., Wang, R., Zhou, Y., Fei, L., Sun, H., Lai, S., Saadatpour, A., Zhou, Z., Chen, H., Ye, F., et al. (2018). Mapping the Mouse Cell Atlas by Microwell-Seq. Cell 172, 1091–1107.e1017. 10.1016/j.cell.2018.02.001.

Jacob, A., Morley, M., Hawkins, F., McCauley, K.B., Jean, J.C., Heins, H., Na, C.L., Weaver, T.E., Vedaie, M., Hurley, K., et al. (2017). Differentiation of Human Pluripotent Stem Cells into Functional Lung Alveolar Epithelial Cells. Cell stem cell 21, 472–488 e410. 10.1016/j.stem.2017.08.014.

Juul, N.H., Stockman, C.A., and Desai, T.J. (2020). Niche Cells and Signals that Regulate Lung Alveolar Stem Cells In Vivo (Mar, 10.1101/cshperspect.a035717, 2020). Cold Spring Harbor perspectives in biology 12. ARTN a040303 10.1101/cshperspect.a040303.

Katzen, J., and Beers, M.F. (2020). Contributions of alveolar epithelial cell quality control to pulmonary fibrosis. Journal of Clinical Investigation 130, 5088–5099. 10.1172/Jci139519.

Kumar, N., Arthur, C.P., Ciferri, C., and Matsumoto, M.L. (2020). Structure of the secretory immunoglobulin A core. Science 367, 1008-+. 10.1126/science.aaz5807.

Lederer, D.J., and Martinez, F.J. (2018). Idiopathic Pulmonary Fibrosis. New England Journal of Medicine 378, 1811–1823. 10.1056/Nejmra1705751.

Liew, P.X., and Kubes, P. (2019). The Neutrophil’s Role during Health and Disease. Physiol Rev 99, 1223–1248. 10.1152/physrev.00012.2018.

Liu, H.J., Fang, S.C., Wang, W., Cheng, Y.S., Zhang, Y.M., Liao, H., Yao, H.H., and Chao, J. (2016). Macrophage-derived MCPIP1 mediates silica-induced pulmonary fibrosis via autophagy. Particle and fibre toxicology 13. ARTN 55 10.1186/s12989-016-0167-z.

Liu, Q.Z., Liu, K., Cui, G.Z., Huang, X.Z., Yao, S., Guo, W.K., Qin, Z., Li, Y., Yang, R., Pu, W.J., et al. (2019). Lung regeneration by multipotent stem cells residing at the bronchioalveolar-duct junction (vol 51, pg 728, 2019). Nature genetics 51, 766–766. 10.1038/s41588-019-0388-9.

Mcgowan, S.E. (1992). Extracellular-Matrix and the Regulation of Lung Development and Repair. Faseb Journal 6, 2895–2904.

Naizhen, X., Kido, T., Yokoyama, S., Linnoila, R.I., and Kimura, S. (2019). Spatiotemporal Expression of Three Secretoglobin Proteins, SCGB1A1, SCGB3A1, and SCGB3A2, in Mouse Airway Epithelia. Journal of Histochemistry & Cytochemistry 67, 453–463. 10.1369/0022155419829050.

Noble, P.W., Barkauskas, C.E., and Jiang, D.H. (2012). Pulmonary fibrosis: patterns and Perpetrators. Journal of Clinical Investigation 122, 2756–2762. 10.1172/JCI60323.

Parimon, T., Yao, C.F., Stripp, B.R., Noble, P.W., and Chen, P. (2020). Alveolar Epithelial Type II Cells as Drivers of Lung Fibrosis in Idiopathic Pulmonary Fibrosis. International journal of molecular sciences 21. Artn 2269 10.3390/Ijms21072269.

Qiu, X.Y., Zhu, X.H., Zhang, L., Mao, Y.T., Zhang, J., Hao, P., Li, G.H., Lv, P., Li, Z.X., Sun, X., et al. (2003). Human epithelial cancers secrete immunoglobulin G with unidentified specificity to promote growth and survival of tumor cells. Cancer research 63, 6488–6495.

Reynolds, S.D., Reynolds, P.R., Pryhuber, G.S., Finder, J.D., and Stripp, B.R. (2002). Secretoglobins SCGB3A1 and SCGB3A2 Define Secretory Cell Subsets in Mouse and Human Airways. American journal of respiratory and critical care medicine 166, 1498–1509. 10.1164/rccm.200204-285OC.

Snoeck, V., Peters, I.R., and Cox, E. (2006). The IgA system: a comparison of structure and function in different species. Vet Res 37, 455–467. 10.1051/vetres:2006010.

Tan, S., and Chen, S. (2021). Macrophage Autophagy and Silicosis: Current Perspective and Latest Insights. International journal of molecular sciences 22, 453. 10.3390/ijms22010453.

Thannickal, V.J., Toews, G.B., White, E.S., Lynch, J.P., and Martinez, F.J. (2004). Mechanisms of pulmonary fibrosis. Annu Rev Med 55, 395–417. 10.1146/annurev.med.55.091902.103810.

Theocharis, A.D., Skandalis, S.S., Gialeli, C., and Karamanos, N.K. (2016). Extracellular matrix structure. Adv Drug Deliver Rev 97, 4–27. 10.1016/j.addr.2015.11.001.

Turnerstokes, L., Haslam, P., Jones, M., Dudeney, C., Lepage, S., and Isenberg, D. (1990). Autoantibody and Idiotype Profile of Lung Involvement in Autoimmune Rheumatic Disease. Ann Rheum Dis 49, 160–162. Doi 10.1136/Ard.49.3.160.

Wells, A.U., Brown, K.K., Flaherty, K.R., Kolb, M., and Thannickal, Victor J. (2018). What’s in a name? That which we call IPF, by any other name would act the same. European Respiratory Journal 51, 1800692. 10.1183/13993003.00692-2018.

Yamamoto, Y., Gotoh, S., Korogi, Y., Seki, M., Konishi, S., Ikeo, S., Sone, N., Nagasaki, T., Matsumoto, H., Muro, S., et al. (2017). Long-term expansion of alveolar stem cells derived from human iPS cells in organoids. Nature methods 14, 1097-+. 10.1038/Nmeth.4448.

Yao, C.F., Guan, X.R., Carraro, G., Parimon, T., Liu, X., Huang, G.L., Mulay, A., Soukiasian, H.J., David, G., Weigt, S.S., et al. (2021). Senescence of Alveolar Type 2 Cells Drives Progressive Pulmonary Fibrosis. American journal of respiratory and critical care medicine 203, 707–717. 10.1164/rccm.202004-1274OC.

Zhang, X.X., Lan, Y.J., Xu, J.Y., Quan, F., Zhao, E.J., Deng, C.Y., Luo, T., Xu, L.W., Liao, G.M., Yan, M., et al. (2019). CellMarker: a manually curated resource of cell markers in human and mouse. Nucleic acids research 47, D721–D728. 10.1093/nar/gky900.

Zhou, Z., Jiang, R., Yang, X., Guo, H., Fang, S., Zhang, Y., Cheng, Y., Wang, J., Yao, H., and Chao, J. (2018). circRNA Mediates Silica-Induced Macrophage Activation Via HECTD1/ZC3H12A-Dependent Ubiquitination. Theranostics 8, 575–592. 10.7150/thno.21648.

